# Direction and orientation preferences in mouse superior colliculus and its retinal inputs align with topographic axes atop locally mixed tuning

**DOI:** 10.64898/2025.12.21.695778

**Authors:** Zhewen He, María Florencia González Fleitas, Raikhangul Gabdrashova, Sylvia Schröder

**Affiliations:** Institute of Ophthalmology, University College London, London WC1E 6BT, UK; School of Life Sciences, University of Sussex, Brighton BN1 9QG, UK

## Abstract

In mouse superior colliculus (SC), neurons are robustly tuned to motion direction and orientation, but it remains controversial whether these preferences are spatially organized, and how this organization relates to that found in the retinal input to the SC. We addressed these questions by combining two-photon calcium imaging of retinal boutons and SC neurons in superficial SC with Neuropixels recordings across the full depth of SC. We then asked how direction and orientation preferences depend on visual-field location and how tuning similarity among units depends on lateral and vertical distance in the SC. Retinal boutons were strongly tuned, and their preferences closely matched the retinal topographic organization of four motion directions and orientations previously described in the retina, rather than a global map in which a single direction or orientation dominates each visual-field location. With increasing depth, SC neurons progressively deviated from this organization. Superimposed on the identified topography, local clustering of tuning preferences in retinal boutons and SC neurons was weak and confined to very small spatial scales. Together, these results show that retinal inputs and neurons in mouse SC represent multiple directions and orientations for each location in the visual field, likely supporting flexible readout for diverse visually guided behaviors.

## Introduction

Visual feature maps in the cortex can range from smoothly organized columns to largely disordered “salt-and-pepper” arrangements. In carnivores and primates, orientation and direction preferences in primary visual cortex (V1) form orderly maps (Basole et al., 2003; Hubel and Wiesel, 1968, 1962; Ibbotson and Jung, 2020), whereas in rodent V1, similarly tuned neurons are embedded in a largely salt-and-pepper background with only weak local clustering (Kaschube, 2014; Ohki et al., 2005). While the SC is the major retinorecipient structure in the mouse midbrain and plays a central role in visually-guided orienting, escape, and prey capture (Hoy and Farrow, 2025; Wheatcroft et al., 2022), its functional organization remains debated (Ahmadlou and Heimel, 2015; Chen et al., 2021; De Malmazet et al., 2018; Feinberg and Meister, 2014; Inayat et al., 2015; Kasai and Isa, 2021; Li et al., 2020). Neurons in superficial SC (sSC) are robustly tuned to direction and orientation (De Franceschi and Solomon, 2018; Inayat et al., 2015; Ito et al., 2017; Wang et al., 2010), but it is unclear whether their preferences form stereotyped maps across the visual field, are locally mixed, or follow another form of spatial organization.

Previous studies have reported conflicting organizations of direction and orientation in mouse SC. Some found strong clustering and global structure (Feinberg and Meister, 2014; Li et al., 2020), including concentric orientation maps centered on the nose (Ahmadlou and Heimel, 2015; De Malmazet et al., 2018) and a predominant motion direction that reverses near the binocular border (De Malmazet et al., 2018). Other work reported relatively weak or no direction and orientation clustering (Chen et al., 2021; Inayat et al., 2015; Kasai and Isa, 2021). These discrepancies suggest that the presence, strength, and spatial scale of direction and orientation maps in SC may depend on specific experimental conditions.

Importantly, the possible organizations of SC tuning are not limited to a binary distinction between salt-and-pepper tuning and a global map in which each visual-field location is dominated by a single direction or orientation preference. Work in the retina has revealed a different form of organization: direction-selective and orientation-selective ganglion cells (DSGCs and OSGCs) align their preferred directions and orientations with multiple position-dependent lines in spherical visual coordinates (Laniado et al., 2025; Sabbah et al., 2017; Tiriac et al., 2022; Vita et al., 2024). DSGCs form four main classes whose preferred motion directions align with longitudinal lines related to optic flow during forward and backward translation and upward and downward motion (Sabbah et al., 2017). OSGCs form an analogous topography, in which preferred orientations align with longitudinal or latitudinal lines defined by multiple poles (Laniado et al., 2025). We refer to these resulting location-dependent model predictions as topographic preferences. Because most retinal ganglion cell (RGC) types, including all direction-selective types, project retinotopically to sSC (Ellis et al., 2016; Kay et al., 2011), this organization provides a concrete alternative to both salt-and-pepper tuning and single-preference maps: each visual-field location may receive input encoding four direction and four orientation preferences.

This distinction between forms of global organization is functionally important. If local SC populations were dominated by only one direction or orientation at each visual-field location, information about other directions or orientations would be weak or absent locally, limiting the ability to detect behaviorally relevant motion or edges in that part of the visual field. Conversely, if multiple directions and orientations are represented locally, SC could preserve broad feature coverage at each location.

Here, we combined two-photon calcium imaging of retinal boutons and SC neurons in posterior sSC with Neuropixels recordings across the full depth of anterior SC. Using receptive-field (RF) mapping, we positioned each unit in visual space and compared its preferred direction and orientation with two sets of predictions expected to be consistent across individual animals: those of the retinal topographic model, which predicts four direction and four orientation preferences at each visual-field location, and those of a single-preference map, in which one direction or orientation dominates each location. We then quantified local clustering and examined how alignment to the topographic preferences changed from retinal boutons to SC neurons and with depth. Our results show that retinal inputs to SC closely match the topography of multiple direction and orientation preferences described in the retina, whereas SC neurons progressively deviate from this organization and exhibit locally mixed tuning with weak clustering.

## Results

### Retinal boutons and sSC neurons are robustly tuned to motion direction and orientation with similar cardinal biases

To compare the functional organization in retinal inputs and SC neurons, we imaged calcium signals in synaptic boutons of RGC axons and in neurons in the posterior right SC. In one group of mice, we expressed a synaptically targeted calcium indicator (Dreosti et al., 2009) in RGCs by intravitreal injection of AAV2-SyGCaMP6f into the left eye (Figure 1A). In another group of mice, we expressed a cytosolic calcium indicator (Chen et al., 2013) in neurons in the superficial layers of the right SC by injecting AAV1-GCaMP6f (Figure 1C). To expose the posterior SC representing the top-left peripheral visual field, we used a cylindrical cranial implant to displace the transverse sinus without damaging cortex. During two-photon imaging, mice were awake and head-fixed on a treadmill, allowing them to run freely. Recorded movies of calcium signals were motion-corrected and segmented into regions of interest (ROIs) with an adapted version of Suite2p (Pachitariu et al., 2016) that accounted for lateral and axial brain motion.

**Figure 1.**
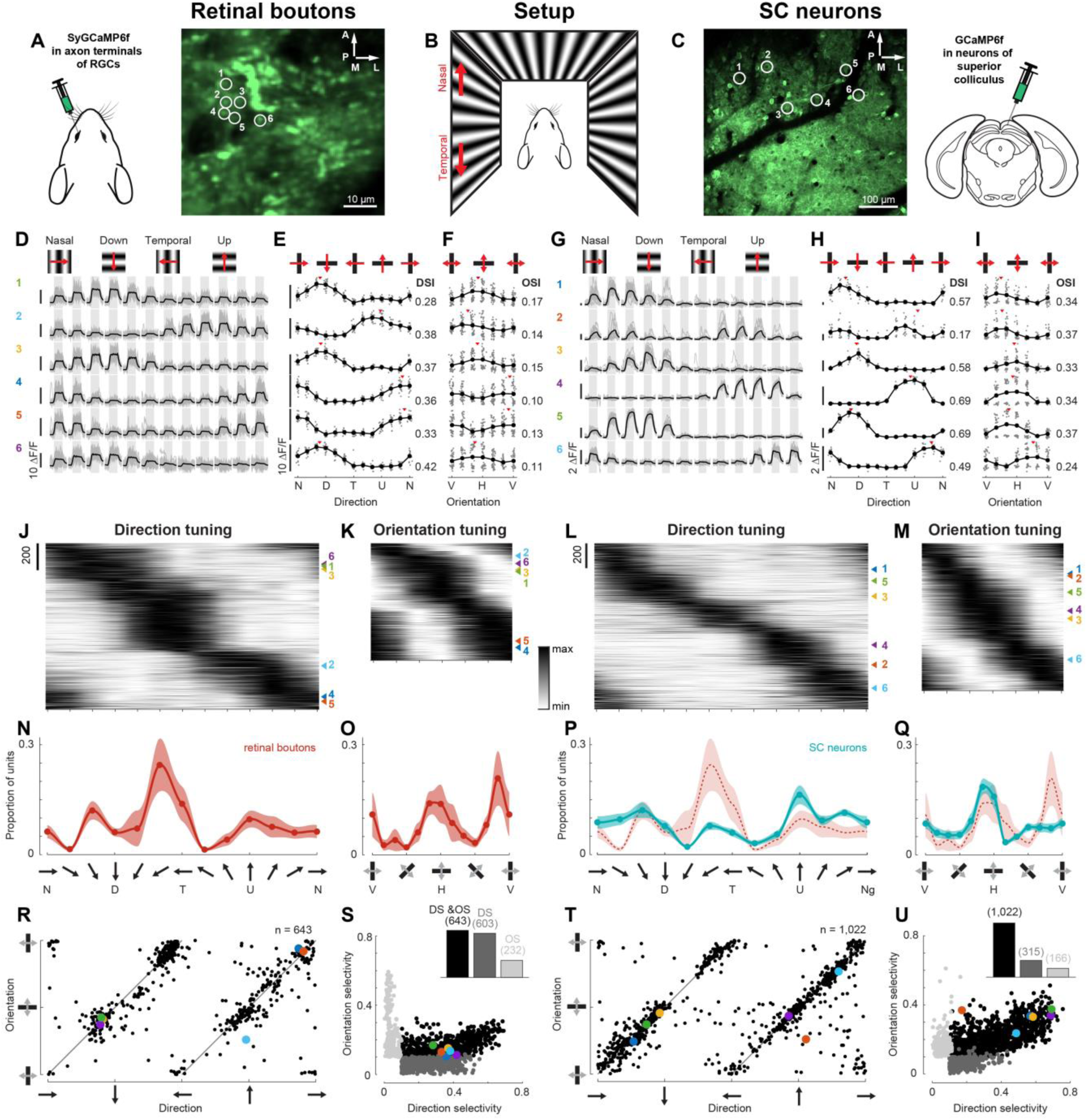
Retinal boutons and neurons in the SC were well tuned to motion direction and orientation. A. AAV-SyGCaMP6f was delivered to RGCs via intravitreal injection (left). Example average image of retinal boutons in the SC (right). B. Visual stimuli were presented on three monitors covering the visual field from −135° to 135° azimuth and −42° to 42° elevation. Grating stimuli covered all screens. Stimulus directions—specifically nasal and temporal—were described from the perspective of the left eye, because RFs of all recorded units were in the left hemifield. C. AAV-GCaMP6f is delivered to SC neurons via injection (right). Example average image of neurons in the SC (left). D. Single-trial fluorescence traces (gray) and mean fitted traces (black) of example boutons (marked in A) in response to sinusoidal gratings drifting in 12 directions. Gray shading indicates the 2 s stimulus presentations. E–F. Direction (E) and orientation (F) tuning curves of example boutons (same as in D), showing single-trial response amplitudes (gray dots), mean amplitudes (black dots), and preferred direction/orientation (red triangles). N – nasal, D – down, T – temporal, U – up, V – vertical, H – horizontal. G–I. As in D–F, for example SC neurons (marked in C). J–M. Direction (J) and orientation (K) tuning curves of all DS and OS retinal boutons recorded across all sessions, sorted by their preferred direction of motion or orientation. Each tuning curve was scaled to a range between 0 and 1. Triangles indicate tuning curves of example boutons from A. Corresponding data for SC neurons in L and M. N–Q. Mean distribution (± SEM) of preferred directions (N) and orientations (O) across 13 and 12 sessions of retinal boutons (red). Datasets with fewer than 20 selective units were excluded. Corresponding data for SC neurons (teal) in P and Q (12 sessions each). For comparison, mean distributions of retinal boutons are overlaid (red; same data as in N, O). R. Preferred direction versus orientation for retinal boutons selective to both features. Example boutons from A marked in color. S. Direction versus orientation selectivity for all tuned retinal boutons. Example boutons from A marked in color. Insets show histograms of selectivity types. T, U. As in R and S, for SC neurons.

We measured tuning to direction of motion and orientation of drifting sinusoidal gratings. Visual stimuli were presented on three monitors surrounding the animal, covering −135° to 135° azimuth and −42° to 42° elevation (Figure 1B). All stimuli were first defined in spherical visual coordinates and then mapped onto the flat monitor surfaces. Gratings moved in 12 directions (30° steps) at a fixed spatial frequency of 0.08 cycles/°, temporal frequency of 2 cycles/s, and 100% contrast. For each bouton and neuron, we fitted responses with a single temporal kernel aligned to stimulus onset and an independent amplitude per trial, separating responses to the current stimulus from residual activity induced by the previous stimulus. This model captured responses well (Figure 1D,G; mean ± standard deviation (SD) explained variance 0.32 ± 0.20 for boutons, 0.44 ± 0.23 for neurons), and identified 65% of retinal boutons (2,391/3,677) and 50% of SC neurons (2,378/4,769) as responsive to drifting gratings. Based on the fitted amplitudes, we calculated the vector average over all motion directions and all orientations. For both direction and orientation, preference and selectivity were defined as the angle and length of the resulting vector (Figure 1E,F,H,I). This measure of orientation tuning has also been termed axis tuning, because it is based on responses to motion along an axis (opposite directions combined) rather than to static stimuli. Orientation preferences were consistent across moving and static stimuli (Figure S1K,L), so we will use the more common term *orientation selectivity*. A bouton or neuron was considered direction or orientation tuned if its selectivity was significantly larger than zero and had a value of at least 0.1. Direction and orientation tuning were not treated as mutually exclusive, because our analyses ask how much information about each stimulus feature is present in the response, rather than assigning units to exclusive functional classes.

Retinal boutons and SC neurons were frequently selective to direction or orientation and exhibited a shared bias toward cardinal directions and orientations. Among responsive boutons, 25% were direction-selective (DS) only, 10% orientation-selective (OS) only, 27% selective to both features, and 38% non-selective (Figure 1S). Responsive SC neurons comprised a larger fraction of units selective to both direction and orientation (43%), fewer units selective to a single feature (DS-only: 13%; OS-only: 7%), and a similar non-selective fraction (37%) (Figure 1U). In a subset of SC neurons, we also presented moving bars and static gratings and found that preferred directions and orientations were highly consistent, confirming that tuning was largely invariant to stimulus class (Figure S1A–L) (Ahmadlou and Heimel, 2015; Inayat et al., 2015). Across both retinal boutons and SC neurons, preferred directions and orientations spanned the full range, but were overrepresented near cardinal angles (0°, 90°, 180°, 270°; Figure 1J–M), which was consistent across recordings (Figure 1N–Q) and in line with previous reports in RGCs (Baden et al., 2016; Kerschensteiner and Hardesty, 2022; Laniado et al., 2025; Nath and Schwartz, 2016; Sabbah et al., 2017; Tiriac et al., 2022; Vita et al., 2024) and SC neurons (Chen et al., 2021; Inayat et al., 2015). Preferred directions strongly predicted preferred orientations (Figure 1R,T), and median direction selectivity exceeded orientation selectivity (boutons: 0.28 vs 0.16; neurons: 0.29 vs 0.19; Figure 1S,U, Figure S1M,N).

A direct comparison between retinal boutons and SC neurons revealed subtle differences in tuning. Both boutons and neurons showed cardinal biases, but boutons displayed a stronger bias toward temporal motion (consistent with optic flow during forward locomotion), while SC neurons were more biased toward near-horizontal orientations (Figure 1N–Q). Median direction selectivity was similar across populations (0.28 for boutons vs 0.29 for neurons; p = 0.61, Wilcoxon rank-sum test; Figure S1M), whereas median orientation selectivity was significantly higher in neurons (0.16 vs 0.19; p = 0.004, Wilcoxon rank-sum test; Figure S1N). Together, these results show that retinal boutons and SC neurons in posterior SC share robust direction and orientation tuning with cardinal biases, but differ in the balance between directional bias and orientation selectivity.

### Retinal topography of direction and orientation preferences is reflected in superficial SC

Elegant studies in mouse retina have shown that tuning preferences of DSGCs and OSGCs are systematically organized in retina-centered spherical coordinates (Laniado et al., 2025; Sabbah et al., 2017). DSGCs form four main classes whose preferred motion directions at each retinal location align with longitudinal lines defined by the body axis and the gravitational axis, corresponding to optic flow during forward, backward, upward and downward self-motion (Sabbah et al., 2017) (Figure 3A). OSGCs form an analogous spherical topography, in which preferred orientations align either with one of two longitudinal fields or two latitudinal fields, that is, concentric lines around one of two axes (Laniado et al., 2025) (Figure 3B,C). These retinal studies thus provide a geometric model that predicts, for any given RF position in visual space, which motion directions and edge orientations are preferred by RGCs.

To relate our SC data to this topographic organization of the retina, we first mapped RFs of retinal boutons and SC neurons using sparse noise (Figure 2A–F), which yielded valid RF fits for the majority of direction- or orientation-tuned units (boutons: 85% and 86%; neurons: 66% and 67%). To verify that small eye movements had not biased these estimates, we re-fitted RF positions in a subset of sessions for which we recorded pupil position, simultaneously estimating a linear transform from pupil displacement to gaze shift in visual degrees (Figure S2.1A–D). RF center positions were minimally affected: the median Euclidean distance between corrected and uncorrected centers was 0.55° for retinal boutons and 0.71° for SC neurons (Figure S2.1E–H), well below the typical RF diameter of ∼10°. We therefore used uncorrected RF fits for subsequent analyses.

**Figure 2.**
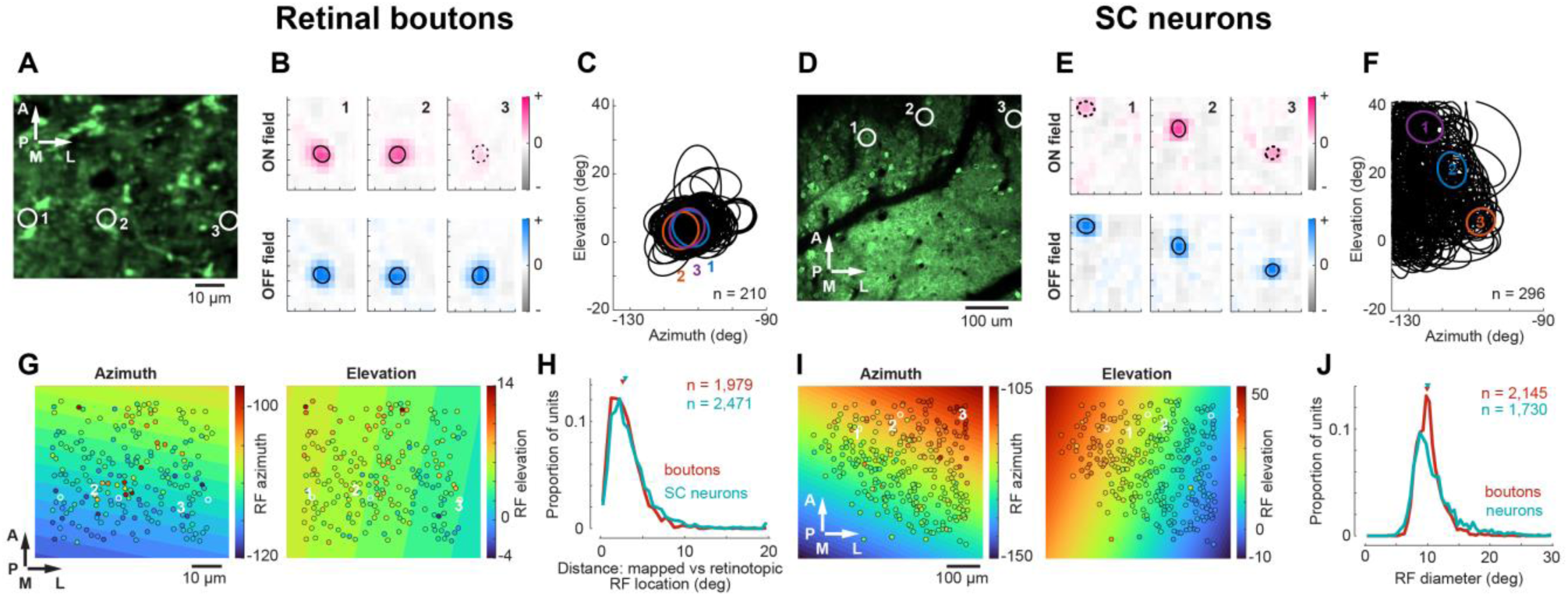
RF locations deviated from retinotopy by about one quarter of their diameter. **A.** Example average image of retinal boutons in the SC. **B.** ON (top) and OFF (bottom) receptive subfields of 3 example retinal boutons (marked in A). Outlines (center ± 1 SD of fitted 2D Gaussian) are shown in black. Examples 1 and 2 had significant ON and OFF subfields, while example 3 had only an OFF subfield (dashed outline of the ON subfield is shown for reference). **C.** RF outlines for all retinal boutons recorded in session from A. Example boutons marked in color. D–F. Same as in A–C, for an example recording of SC neurons (same FOV as in Figure 1C). **G.** Relationship between position in the FOV and RF azimuth (left) or elevation (right) for retinal boutons from the session in A. The colored background gradients represent the best linear fit between FOV and RF positions. **H.** Distances between mapped and retinotopic (predicted from FOV position) RF locations for retinal boutons (red) and SC neurons (teal). Retinal boutons had slightly smaller distances than SC neurons (medians (triangles): 2.6° and 2.9° for boutons and SC neurons; p < 1e-6, Wilcoxon rank-sum test). **I.** Same as in G, for SC neurons and example session from D. **J.** RF diameters (2 SDs of the fitted Gaussian) across retinal boutons (red) and SC neurons (teal). RFs were fitted while correcting for eye movements (see Methods), which modestly affected RF size but not RF position. SC neurons had slightly but significantly larger RF diameters than retinal boutons (medians (triangles): 9.9° and 10.0° for boutons and SC neurons; p = 0.002, Wilcoxon rank-sum test).

Because RFs could not be mapped for every direction- or orientation-tuned unit, we next asked how well RF locations in each imaging field of view (FOV) could be described by a local retinotopic map. For each FOV, we fitted a simple linear retinotopic map that optimally translated, scaled, and rotated the positions of units in the FOV to best match their RF locations in visual space. The fitted retinotopic maps closely predicted the directly mapped RF positions, with average deviations of 2.63° for boutons and 2.93° for neurons, corresponding to roughly one quarter of the RF diameter (Figure 2G–J). For SC neurons, this allowed us to infer visual-field locations for tuned neurons for which RFs could not be mapped directly. For retinal boutons, however, directly mapped RFs covered a more restricted visual-field range, making the retinotopic fits less well constrained in some datasets. Subsequent analyses therefore used inferred retinotopic RF locations for SC neurons, but only directly fitted RF locations for retinal boutons. The proximity of RFs to monitor edges did not systematically affect measured direction or orientation preferences (Liang et al., 2023; Roth et al., 2018) (Figure S2.2). Nevertheless, we excluded units with RF centers closer than 5° to any monitor edge from further analyses.

The retinal topographic organization of directions and orientations was largely preserved in retinal boutons and partially preserved in SC neurons. For each unit, we projected its RF location onto the axes of the retinal geometric model (Figure 3A–C) and calculated the angular difference between its measured preference and the nearest predicted vector (Figure 3D–G). We used the median angular difference as the effect-size measure of model performance and compared it with two controls: a permutation control that preserved the overall preference distribution but disrupted its relationship with RF location, and a control in which preferences were drawn from a uniform distribution of angles. In retinal boutons, direction preferences differed from the nearest predicted vectors by a median of 9.5°, smaller than expected under the uniform model (20.9–24.2°; Figure 3D,H) or after permuting RF locations (12.3–13.7°; Figure 3H). This comparison shows that direction preferences varied systematically with RF position, despite the small variation in model predictions across the sampled visual-field region, which makes RF permutation a stringent test of position-dependent alignment. Orientation preferences in boutons showed a similar pattern, with a median difference of 7.6°, smaller than both the permuted (9.5–11.0°) and uniform (13.8–17.0°) controls (Figure 3E,I).

**Figure 3.**
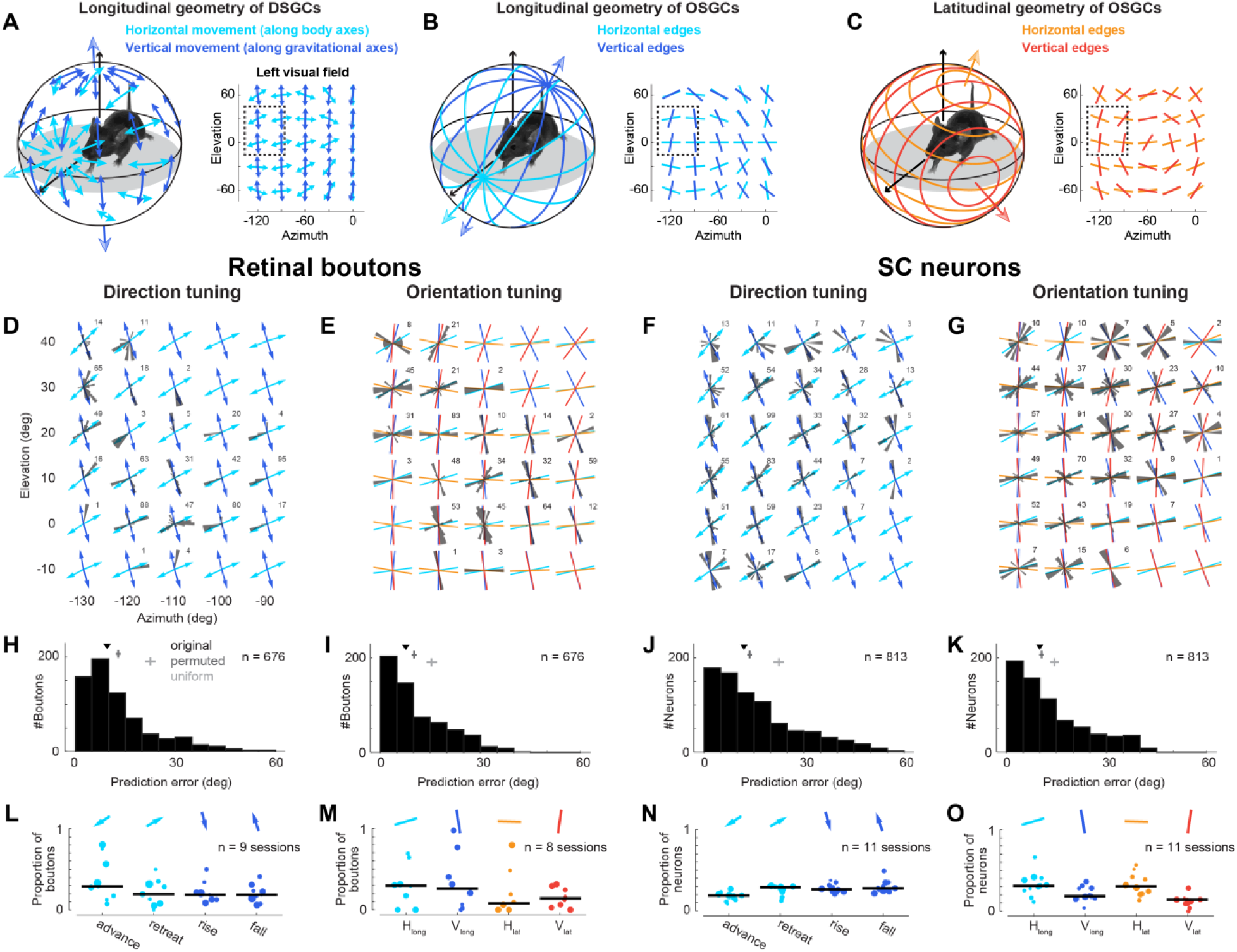
Topography of directions and orientations was largely consistent with topography originally identified in the retina. A–C. Cartoons illustrating the topography of direction preferences (A) of DSGCs and orientation preferences (B, C) of OSGCs for the left eye. Direction preferences align with two axes defining longitudes (A), while orientation preferences align with two axes defining longitudes similar to those for direction (B) and two further axes defining latitudes (C). Dotted rectangle within left visual field marks the part of the visual field that contains RFs of our data (shown in D–G). D–G. Direction (D) and orientation (E) preferences of retinal boutons across visual space. Arrows in D point into directions defined by the DSGCs longitudes (A). Colored lines in E follow orientations of longitudes (B) and latitudes (C) for OSGCs. Numbers indicate unit count in each polar histogram. Corresponding data for SC neurons in F and G. H–K. Angular differences between preferences and closest predicted vectors for direction (H) and orientation (I) across all tuned retinal boutons. The median difference (triangle) of the original data is compared to the median (2.5–97.5 percentile intervals) expected when RF locations were randomly permuted (dark gray) or when preferences were randomly drawn from a uniform distribution (light gray). Direction: 9.55° original, 12.3°–13.7° permuted, 20.9°–24.2° uniform; orientation: 7.6° original, 9.5°–11.0° permuted, 13.8°–17.0° uniform. Corresponding data for SC neurons in N and O. Direction: 12.0° original, 13.0°–14.4° permuted, 20.7°–24.1° uniform; orientation: 10.0° original, 9.8°–11.2° permuted, 13.0°–15.8° uniform. L–O. Classification of all DS (L) and OS (M) boutons according to the closest predicted vector from the geometric model. Each dot shows one recording session (dot size reflects total number of boutons); black line indicates median proportion across sessions. Naming follows previous conventions, where DSGC classes were named after the behavior that elicits their responses, e.g., “advance cells” for DSGCs preferring movement in temporal direction. Arrows and lines at the top are example vectors from a central location in D and E. Corresponding data for SC neurons in N and O.

Direction preferences of SC neurons also followed the geometric predictions, with a median difference of 12.0°, smaller than in permuted (13.0–14.4°) and uniform (20.7–24.1°) controls (Figure 3F,J). However, SC neurons differed more from the predicted directions than retinal boutons (p < 1e-4, Wilcoxon rank-sum test), indicating that the retinal topography was already weakened at the level of sSC neurons. This difference was more pronounced for orientation preferences. In SC neurons, the median angular difference of 10.0° was comparable to the permuted control (9.8–11.2°), although still smaller than the uniform model (13.0–15.8°; Figure 3G,K), and significantly larger than in retinal boutons (p = 1e-4, Wilcoxon rank-sum test). Restricting the analysis to DS-only or OS-only units yielded the same qualitative picture (Figure S3). These findings show that retinal boutons in posterior SC closely preserve the retinal topography of direction and orientation tuning, while SC neurons preserved the topography for direction but deviated from that for orientation. Retinal boutons were biased toward temporal motion-preferring cells, whereas SC neurons showed a more uniform distribution across direction classes. We assigned each tuned bouton or neuron to the predicted direction or orientation vector closest to its measured preference. For direction, we used the four optic-flow classes defined by Sabbah et al. (2017), for example “advance cells”, which prefer optic flow in temporal direction elicited by forward self-motion (Figure 3A). Retinal boutons were strongly biased toward the “advance” class (30% of direction-tuned boutons, Figure 3H), consistent with their stronger bias for temporal stimulus motion in the overall direction distribution (Figure 1N) and with the bias found in DSGCs (Sabbah et al., 2017). SC neurons, by contrast, were more evenly distributed across direction classes, with 19–29% of direction-tuned neurons in each class (Figure 3J). For orientation, boutons were roughly balanced between horizontal and vertical classes (38% vs 40%; Figure 3I), while SC neurons were slightly biased toward horizontal orientation classes (62% vs 33%; Figure 3K).

### Superficial SC and its retinal input lack single-preference maps of direction or orientation

Smoothed maps of population tuning across visual space lacked systematic gradients, and only retinal boutons showed local tuning consistency above chance. While the previous analyses have revealed a topography of four direction and four orientation preferences, we next asked whether there are local biases toward single directions or orientations that could form global maps in the sSC. Visualizing preferred directions and orientations at RF locations across all recordings revealed mixtures of preferences at essentially all positions (Figure 4A–D). Averaging preferences within circular patches of 10° diameter to construct smoothed maps across visual space likewise revealed no systematic gradients or concentric patterns in either boutons or neurons (Figure 4E–H). Because patch averages can obscure heterogeneity among individual units, we quantified local consistency as the inverse circular variance of preferences within each patch (Figure 4I–L). Retinal boutons showed substantially higher local consistency than expected from null distributions obtained by permuting RF locations (direction: 0.49 ± 0.001; orientation: 0.60 ± 0.001; mean ± standard error of the mean (SEM); Figure 4I,J,M,N), and in most individual bouton recordings both direction and orientation consistency exceeded the shuffled control (Figure 4Q,R). In SC neurons, by contrast, local consistency was lower (direction: 0.22 ± 0.0003; orientation: 0.36 ± 0.0004; mean ± SEM; Figure 4K,L) and indistinguishable from null distributions for both features (Figure 4O,P). No recording showed elevated direction consistency and only four recordings showed elevated orientation consistency (Figure 4S,T). For both retinal boutons and SC neurons, consistency was not trivially explained by the number of units per patch (Figure S4). The somewhat higher tuning consistency in local populations of retinal boutons may be explained by subsets of retinal boutons originating from the same RGC and therefore sharing identical tuning. More broadly, the absence of a consistent direction and orientation map in sSC and the relatively weak tuning consistency among boutons and neurons contrasts previous findings, which would predict a clear overrepresentation of near-vertical orientation preferences and a mix of temporal and upward direction preferences in the posterior SC (Ahmadlou and Heimel, 2015; De Malmazet et al., 2024, 2018; Feinberg and Meister, 2014; Li et al., 2020); these discrepancies are addressed in the Discussion.

**Figure 4.**
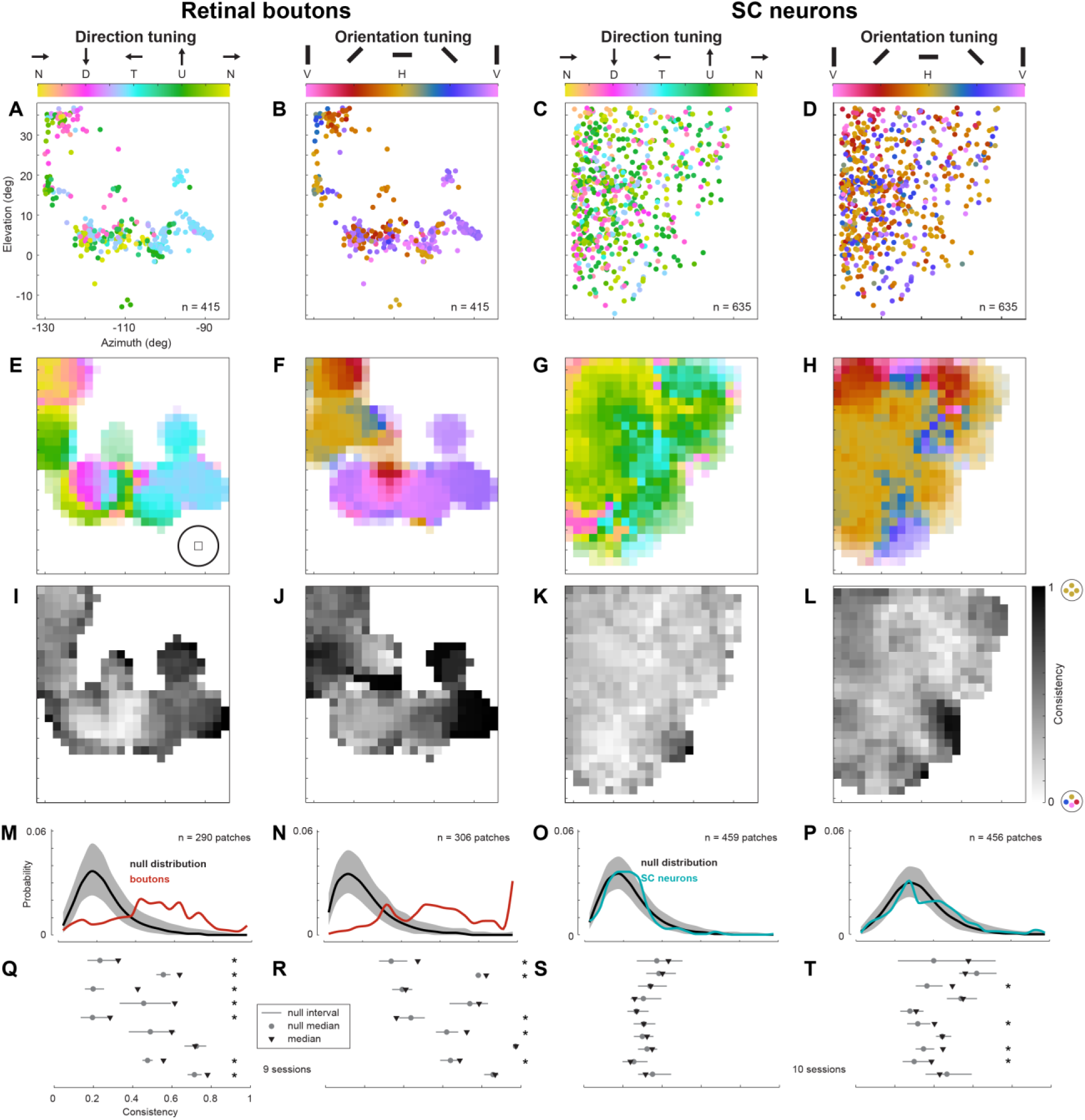
Local populations showed low consistency of direction and orientation preferences. A–D. Direction (A) and orientation (B) preferences for all selective retinal boutons, plotted at their RF locations in visual space. Corresponding data for SC neurons in C and D. E–H. Mean direction (E) and orientation (F) preferences obtained by averaging across boutons within circular patches of 10° diameter in visual space (illustrated by circle in the lower-right corner). Transparency indicates patches with fewer than 20 boutons. Corresponding data for SC neurons in G and H. I–L. Consistency of direction (I) and orientation (J) preferences within 10° patches, quantified as the inverse circular variance of preferences (same pooling as in E and F) for retinal boutons. Corresponding data for SC neurons in K and L. M–P. Distributions of consistency for direction (M) and orientation (N) across patches (red; at least five units per patch) compared to null distribution obtained by permuting RF locations across boutons (black line: median; gray band: 2.5th–97.5th percentile interval). Corresponding data for SC neurons in O and P. Q–T. Mean consistency in direction (Q) and orientation (R) preferences across all patches per recording for retinal boutons (black triangles), compared to corresponding null distribution for each recording (gray; dot: median; vertical line: 2.5th–97.5th percentile interval). Recordings with significant consistency values are marked by a star. Corresponding data for SC neurons in S and T.

### Local functional clustering in sSC is weak and limited to very small spatial scales

Pairs of retinal boutons showed more similar tuning preferences than pairs of SC neurons. In example FOVs, preferences of boutons and neurons showed intermingled directions and orientations without obvious clusters (Figure 5A–D). When we considered all pairs of simultaneously recorded units, pairwise differences were smaller than expected for a uniform distribution of preferences (90° for direction, 45° for orientation) but still large: mean differences were ∼75° for direction and 32° for orientation among boutons, and ∼89° and 42° among neurons (Figure 5E–H).

**Figure 5.**
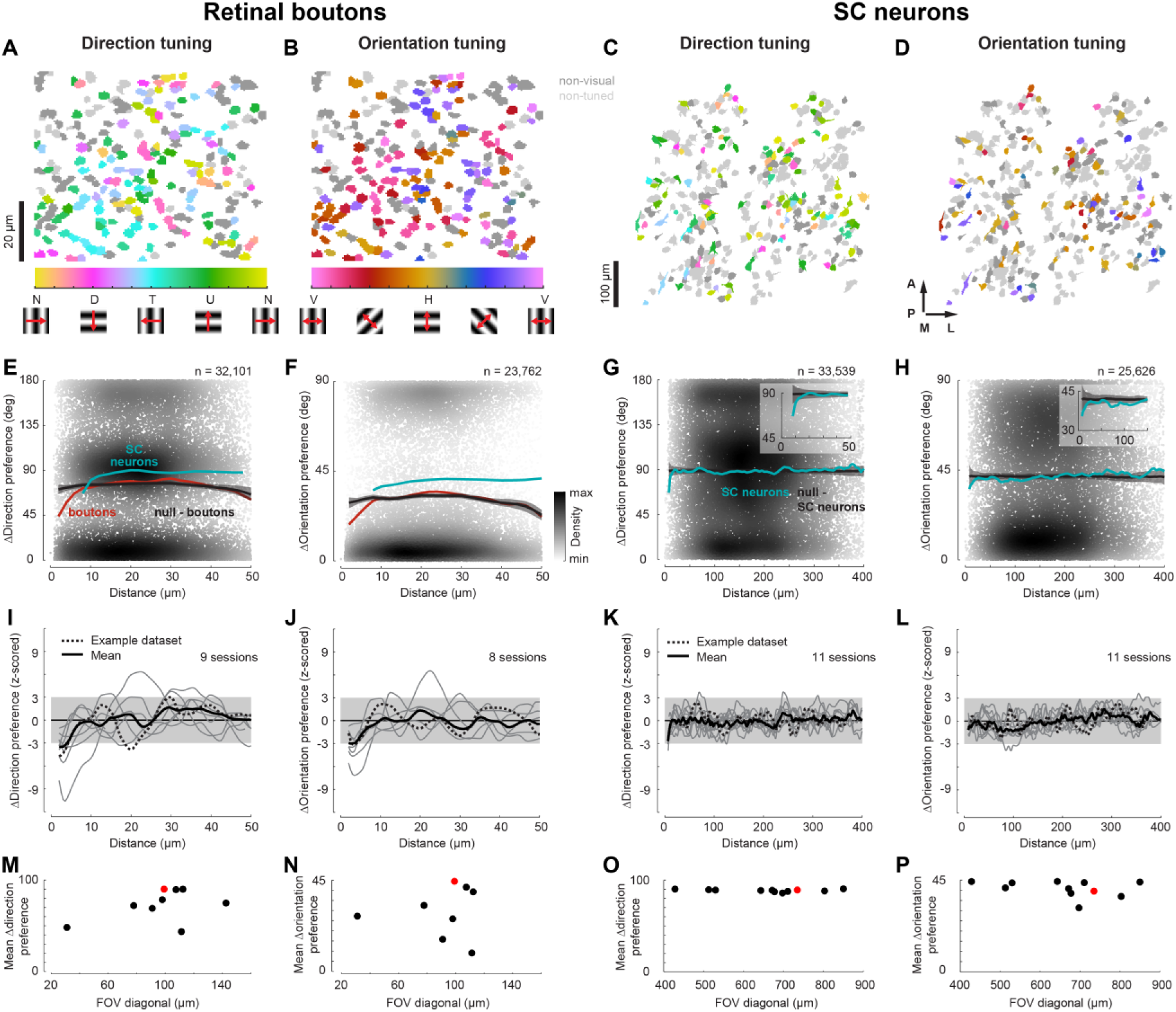
Pairwise similarity of direction and orientation tuning was largely independent of lateral distance. A–D. Examples of direction (A) and orientation (B) preferences of individual retinal boutons in one FOV. Each bouton is plotted at its imaging location; non-responsive boutons are shown in dark gray and non-selective boutons in light gray. Corresponding data for SC neurons in C and D. Neurons from four imaging planes are shown. E–H. Pairwise difference in direction (E) and orientation (F) preferences as a function of lateral distance in the SC (depth ignored) for all pairs of simultaneously recorded retinal boutons (dots colored by density). Mean difference as a function of distance (red) is compared to the null distribution obtained by permuting bouton locations (black line: median; gray band: 2.5th–97.5th percentile interval). For comparison, mean pairwise difference of SC neurons is overlaid (teal; same data as in G, H). Corresponding data for SC neurons in G and H. Insets show zoom-in. I–L. Mean pairwise difference in direction (I) and orientation (J) preference for each recording (gray lines, at least 15 selective units each) as a function of distance, normalized to the null distribution of that recording. Mean (horizontal black line) and ±3 SEMs (gray shade) of the null distribution are shown for comparison. Example recordings from A and B are shown as dotted lines; mean across all recordings depicted as bold black line. Corresponding data for SC neurons in K and L. M–P. Mean pairwise difference in direction (M) and orientation (N) preference for each recording (at least 15 selective units each) as a function of FOV size (length of diagonal). Example from A and B shown in red. Corresponding data for SC neurons in O and P.

Pairwise tuning differences showed only weak dependence on lateral distance and departed from chance levels only for very close pairs. Pairwise differences as a function of lateral distance in the SC closely followed null distributions obtained by randomly permuting unit locations within each FOV (Figure 5E–H). For retinal boutons, only pairs separated by ≤10 μm had direction and orientation preferences that were clearly more similar than expected from the null distribution (Figure 5E,F). For SC neurons, elevated similarity was restricted to pairs within <12 μm, with orientation differences remaining slightly below the null distribution up to ∼130 μm but with a very small absolute deviation (<2°; Figure 5G,H). Across recordings, this weak clustering at very short distances remained detectable for boutons but was rare for neurons (Figure 5I–L). Because bouton recordings covered smaller FOVs than neuron recordings, we tested whether FOV size could explain the higher similarity among boutons. This was not the case. Mean pairwise differences in bouton recordings showed no systematic relationship with FOV size, and some bouton recordings exhibited mean differences as large as those in neuron recordings with FOVs at least four times bigger (Figure 5M–P). This weak clustering persisted when we measured distances in visual space rather than in FOVs, and when we used different visual stimuli. When we considered pairwise differences as a function of RF distance, bouton and neuron pairs showed the same pattern, with deviations from chance confined to RF pairs within a few degrees (Figure S5A–D). Pairwise tuning differences derived from moving bars or static gratings were similar to those for drifting gratings (Figure S5E–G).

### Tuning across SC depth was heterogeneous and weakly related to the retinal topography

Neuropixels recordings across the full depth of the SC revealed robust tuning to motion direction and orientation. We recorded spiking activity using Neuropixels probes inserted vertically through the full depth of the SC in an experimental setup similar to the two-photon imaging experiments (Figure 6A,B). Single units were isolated by automatic spike sorting using Kilosort2.5 (Pachitariu et al., 2024) and quality metrics followed by manual curation. The border between sSC and deep SC (dSC) was determined from visually evoked local field potentials, current source density, and histology (Figure S6A–C). Around 58% of responsive neurons in the SC were well tuned (Figure 6C–E,I,K) and the population was biased towards cardinal directions and orientations (Figure 6F,G). Preferred directions were strongly correlated with preferred orientations (Figure S6D). Direction selectivity in sSC was generally higher than orientation selectivity (Figure 6K), while, across all depths, OS units were more numerous than DS units (Figure S6E).

**Figure 6.**
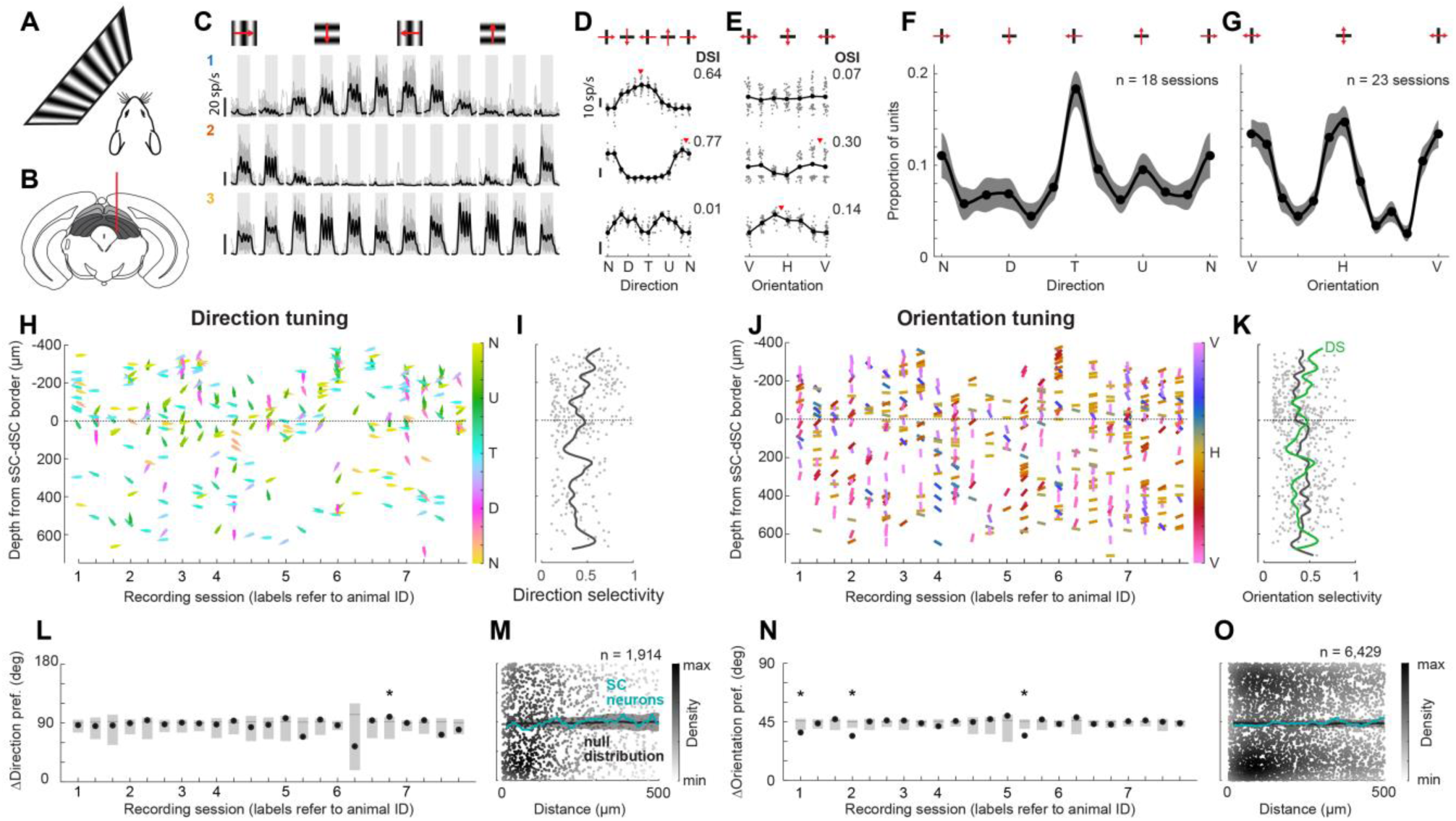
Electrophysiological recordings across SC depth confirmed the lack of functional clustering. A. Visual stimuli were presented on a single monitor covering −95° to 0° azimuth and −27° to 51° elevation. Drifting grating stimuli covered the full screen. **B.** Neuropixels probes were used to record single units from sSC (light gray) and dSC (dark gray). **C.** Single-trial (gray) and mean (black) spike rate traces of example units in response to sinusoidal gratings drifting in 12 directions. Grating presentations (2 s each) are indicated by gray shading. D, E. Direction (D) and orientation (E) tuning curves for example units in C, showing single-trial response amplitudes (gray dots), mean amplitudes (black dots), and preferred directions and orientations if significantly tuned (red triangles). F, G. Mean distribution (± SEM) of preferred directions (F) and orientations (G) across all recording sessions. Datasets with fewer than 10 selective units were excluded. **H.** Preferred direction of all recorded units, as a function of depth in SC for each recording session (multiple sessions per animal). **I.** Direction selectivity of all units as a function of depth in SC. J, K. Same as in H and I, for orientation tuning. For comparison, direction selectivity profile (green) from I is overlaid in K. **L.** Mean of pairwise direction difference of each recording (dots), compared to corresponding null distribution (gray line: median; light-gray box: 2.5th–97.5th percentile interval). **M.** Pairwise difference in direction preference as a function of depth separation in SC for all pairs of simultaneously recorded units (dots colored by density). Mean difference as a function of separation (teal) is compared to null distribution obtained by permuting unit depths (black line: median; gray band: 2.5th–97.5th percentile interval). N, O. Same as in L and M, for orientation tuning.

Direction and orientation preferences showed no consistent pattern with SC depth. When preferred directions and orientations were sorted by depth within each recording, no systematic trend or layering was revealed (Figure 6H,J), even though RFs of simultaneously recorded neurons largely overlapped (Figure 7A,B). Quantitatively, median pairwise tuning differences were close to the values expected for a uniform distribution of preferences (90° for direction, 45° for orientation) and rarely deviated from the medians obtained after randomly permuting units across recordings (Figure 6L,N). Pairwise differences were also independent of distance in depth between neurons: mean tuning differences as a function of depth separation closely followed null distributions based on permuted depths for both direction and orientation (Figure 6M,O).

**Figure 7.**
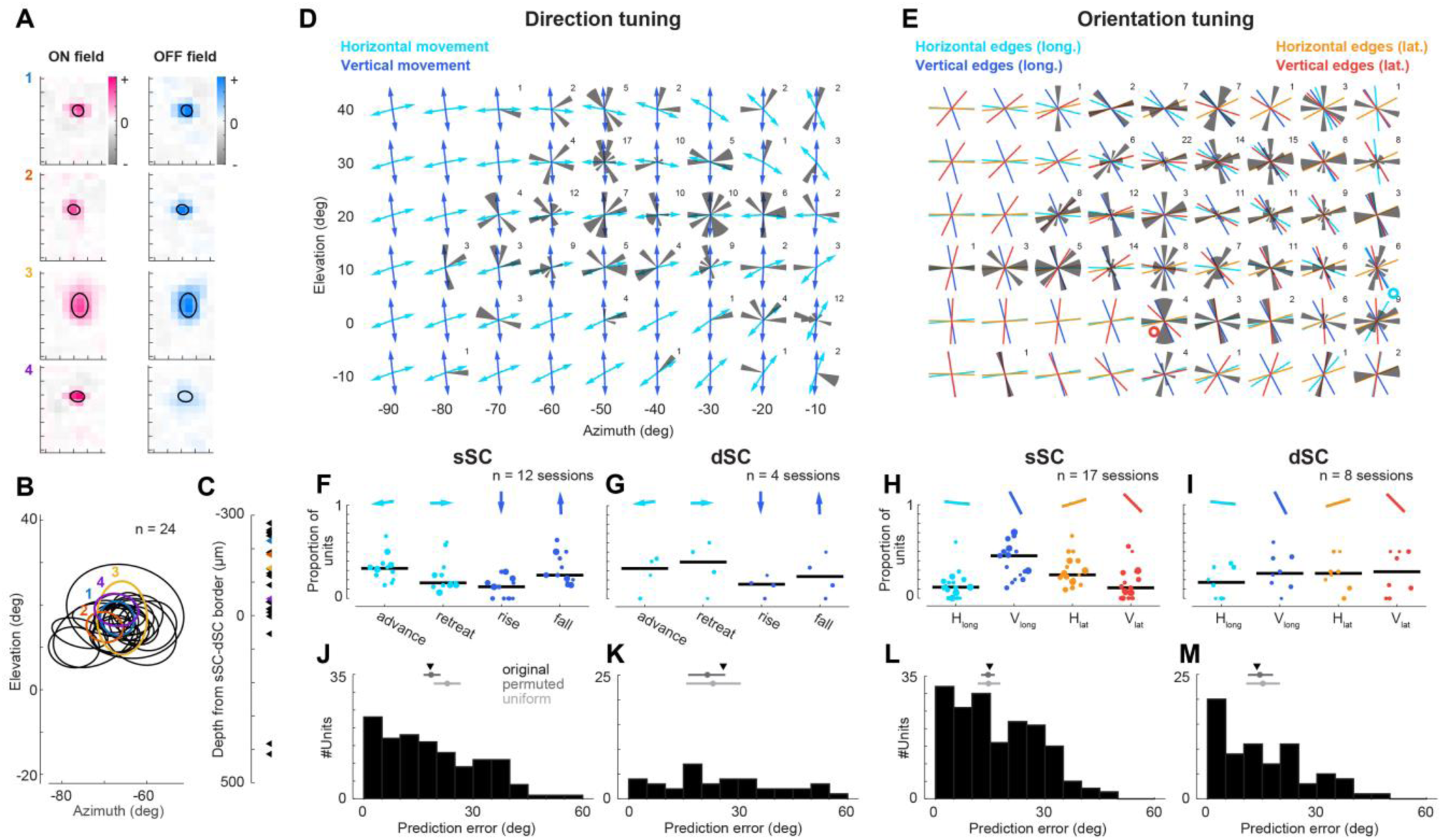
Retinal topography was progressively weakened from superficial to deep SC. A. ON and OFF RFs of four example units. RF outlines (center ± 1 SD of fitted 2D Gaussian) shown in black. **B.** RF outlines of all units recorded in one session, including examples in A (colored). **C.** Depth distribution of units with RFs from B. Depths were aligned to the border between sSC and dSC. D, E. Direction (D) and orientation (E) preferences of SC units across visual space, plotted at their mapped RF locations (not retinotopic). Arrows in D point along directions defined by DSGC longitudes (see Figure 3A). Colored lines in E indicate orientations of longitudinal and latitudinal fields for OSGCs (see Figure 3B,C). Numbers indicate population size in each polar histogram. The depicted region of visual space in E includes two poles of the spherical axes: one for horizontal longitudes (cyan circle) and one for vertical latitudes (red circle). F–I. Classification of all DS units in sSC (F) and dSC (G) according to closest predicted direction vector. Each dot shows one recording session (dot size reflects total number of units); black line indicates median proportion across sessions. Arrows at the top illustrate example predicted vectors from a central location in D. Corresponding data for OS units in H and I. J–M. Angular differences between measured direction preferences and the closest predicted vectors in sSC (J) and dSC (K). The median difference (triangle) is compared to the distribution obtained by permuting RF locations (dark gray: median; band: 2.5th–97.5th percentile interval) and to a uniform preference model (light gray: median; band: 2.5th–97.5th percentile interval). sSC: 18.2° original, 16.4°–21.0° permuted, 19.2°–26.5° uniform; dSC: 25.9° original, 16.3°–26.5° permuted, 15.7°–30.7° uniform. Corresponding data for OS units in L and M. sSC: 14.8° original, 12.5°–16.2° permuted, 11.6°–17.8° uniform; dSC: 13.6° original, 11.1°–18.0° permuted, 10.6°–19.8° uniform.

Direction and orientation tuning across the depth of SC showed little resemblance to the retinal spherical topography. As before, we located RFs of recorded neurons within the visual field (Figure 7A–C). When we compared SC tuning to the geometric model derived from retinal RGCs (Laniado et al., 2025; Sabbah et al., 2017), the match between measured and predicted preferences in sSC was weaker than in retinal boutons (Figure 3) and weakest for dSC neurons (Figure 7D–M). In particular, direction and orientation preferences of dSC neurons showed no alignment with the predicted retinal vectors, which was as weak as expected by uniform distributions of preferences (Figure 7L,M). Together, these results are consistent with increasing integration across diverse retinal inputs as one moves away from the retinal input. We noticed that the distributions of direction and orientation classes recorded in sSC using electrophysiology (Figure 7F,H) appear to be different from the distributions in imaged sSC neurons (Figure 3J,K). This may be explained by the difference in RF locations, differences in recorded depth (imaging is biased to the most superficial sSC), or individual differences across mice.

## Discussion

Our results demonstrate that the topographic organization of direction and orientation tuning previously described in the retina is reflected in retinal input to the mouse SC and is progressively transformed within SC. Using two-photon calcium imaging of retinal boutons and SC neurons in posterior monocular SC, together with Neuropixels recordings across the depth of anterior binocular SC, we found robust tuning and shared cardinal biases in both retinal inputs and neurons. Retinal boutons closely followed geometric models derived from retinal organization, which predict four direction and orientation preferences at each visual-field location based on longitude- and latitude-like lines across visual space (Laniado et al., 2025; Sabbah et al., 2017). The alignment to the geometric model decreased from retinal boutons to superficial neurons and further to deep neurons. Local clustering of tuning was weak in all neural populations and confined to very small spatial scales (10–20 μm), while tuning preferences across visual space lacked smooth gradients and stereotyped global patterns.

### Maps or no maps?

Pairwise differences in preferred direction between SC neurons in our data were largely independent of lateral distance, and consistency of preferences within 100-µm neighborhoods was at chance. These findings agree with some studies (Chen et al., 2021; Inayat et al., 2015), but disagree with reports of local clustering of either direction or orientation preferences (Ahmadlou and Heimel, 2015; De Malmazet et al., 2018; Feinberg and Meister, 2014; Kasai and Isa, 2021; Li et al., 2020). But among cluster- or map-positive studies, the strength of pairwise differences varied markedly: for pairs <100 µm apart, median direction differences were ∼40° in some studies (De Malmazet et al., 2018; Li et al., 2020) versus ∼80° in others (Kasai and Isa, 2021), and median orientation differences ranged from ∼30° (Feinberg and Meister, 2014; Kasai and Isa, 2021) to <10° (De Malmazet et al., 2018). At the level of retinal inputs, nearby boutons were reported to have almost identical orientation preferences and more homogeneous direction tuning than in our data (De Malmazet et al., 2024), whereas we observed clear clustering only within ∼10 µm, plausibly reflecting multiple boutons from single RGCs (Hong et al., 2011).

Inconsistencies among previous reports are even more pronounced at the level of global maps. For direction, different studies emphasized a predominance of upward motion, upward-anterior motion, or opposing posterior/anterior biases in monocular versus binocular SC (Ahmadlou and Heimel, 2015; De Malmazet et al., 2018; Dräger and Hubel, 1975; Li et al., 2020). For orientation, neurons near the center of the visual field were described as horizontally biased or largely non-selective (Ahmadlou and Heimel, 2015; De Malmazet et al., 2018). Proposed map layouts range from concentric orientation maps centered on the nose to ∼30°-wide patches of similar orientation without global structure (Ahmadlou and Heimel, 2015; De Malmazet et al., 2018; Feinberg and Meister, 2014), and large differences across individual animals are evident within single studies (e.g., Figure S3 in De Malmazet et al., 2018; Figure 6C in Li et al., 2020). Against this heterogeneous background, our observation of broadly distributed, weakly clustered preferences suggests that strong, stereotyped maps are unlikely to be a universal organizing principle of SC.

Several factors may contribute to the discrepancies across studies. Some focused on anterior SC and the binocular/monocular border, others on more posterior monocular regions, yet both map-like and salt-and-pepper organizations have been reported in both zones, so retinotopic location alone cannot explain the differences. Visual stimuli varied widely (drifting or static gratings, moving bars, dots), distortions from presenting spherical coordinates on flat screens were not always corrected, and some RF estimates were based on response delays of DS units to moving bars, which can misplace RFs and thus inferred maps (De Malmazet et al., 2024, 2018). Stimulus apertures or monitor edges may also contribute, because RF overlap with stimulus boundaries can induce apparent orientation preferences and, in sSC, stimulus-dependent orientation and direction columns (Borges et al., 2025; Liang et al., 2023; Roth et al., 2018; Yaakov et al., 2025). However, such effects should be strongest for RFs near monitor edges and should induce preferences with a predictable relationship to boundary orientation, for example horizontal orientation preferences near vertical edges. They are therefore unlikely to explain all reported map layouts, including concentric orientation maps or large-scale direction biases toward upward, nasal, or temporal motion. In our data, preferred directions and orientations showed no systematic dependence on RF distance to monitor edges. Most experiments used awake animals, but internal state mainly modulates gain rather than preferred direction or orientation (Chen et al., 2021; Ito et al., 2017; Savier et al., 2019; Schröder et al., 2020), making it an unlikely source of disagreement. Finally, biases in cell-type sampling are plausible if stimuli drive only a minority of neurons, implying that different functional subpopulations were emphasized.

A partial resolution emerges when considering the four direction and orientation preferences predicted by the retinal topographic organization to vary smoothly across visual space (Laniado et al., 2025; Sabbah et al., 2017). This framework does not fall neatly onto a binary distinction between salt-and-pepper organization and single-preference maps. Instead, it predicts a third form of organization: multiple topographic preferences are represented at each visual-field location. The topographic organization encompasses concentric orientation layouts near the front, in addition to other geometries proposed in previous studies. Systematic or random shifts in the relative strengths of these direction and orientation channels across animals, recording methods, or stimulus sets could make one channel dominate over others and thereby produce an apparent single-preference map.

Moreover, some studies analyzed DS and OS populations separately and reported strong structure within, but random mixing between, these groups (De Malmazet et al., 2024, 2018). Given that neurons with both direction and orientation selectivity typically exhibit approximately orthogonal preferences, pooling DS and OS populations and including neurons selective to both features will naturally yield more heterogeneous local preference distributions, with broader local coverage of stimulus directions and orientations rather than dominance of a single preference at each visual-field location.

### Retinal topography was progressively transformed by collicular circuits

Retinal boutons in sSC closely followed the topographic models of DSGCs and OSGCs (Laniado et al., 2025; Sabbah et al., 2017; Tiriac et al., 2022). This is expected because 85–90% of RGCs, including all four cardinal DSGC types, project to SC (Ellis et al., 2016; Kay et al., 2011), their terminals predominantly target the superficial layers that we imaged (Dhande and Huberman, 2014; Hong et al., 2011; Huberman et al., 2009), and their projections follow strict retinotopy (Molotkov et al., 2023; Sibille et al., 2022). Our RF measurements confirmed this tight retinotopic alignment.

In sSC neurons, the topographic organization remained evident for direction but was weaker for orientation. Direction selectivity and preferred direction in SC are largely inherited from DSGCs and amplified by similarly tuned intracollicular inputs (Shi et al., 2017; Sibille et al., 2022), while some DS neurons in SC also receive input from non-DSGCs (Kay and Triplett, 2017). Axis-selective responses may be inherited from OSGCs, or may arise from convergent innervation by oppositely tuned DSGCs or local SC circuits (Kay and Triplett, 2017). Our finding that direction preferences of superficial neurons remained well aligned with the retinal topography, whereas orientation preferences deviated more than in boutons, is therefore consistent with a largely preserved feedforward scaffold for direction combined with mixing of multiple retinal channels and local circuitry that blurs topographic orientation organization at the single-neuron level. The weaker agreement with the topographic organization in Neuropixels-recorded superficial neurons likely reflects undersampling of the most superficial sSC layer, which is preferentially targeted by DSGCs.

In dSC, we found almost no agreement with the retinal topographic organization despite robust tuning. Functional classifications in previous studies identified “RGC-like” neuron types that predominate in sSC and more “SC-specific” types that become more prominent with depth (Lee et al., 2020; Li and Meister, 2023), consistent with increasing transformation away from retinal channels. This transformation in dSC is expected given further functional integration within SC and the convergence of signals from other sources, including cortical and brainstem structures (Cang et al., 2024; Liu et al., 2022; Wheatcroft et al., 2022). Together, these results support a hierarchical view in which a retinally derived topographic organization is preserved in retinal boutons, but increasingly mixed across SC layers.

### Parallels with mouse V1: local mixing rather than single-preference maps

Several aspects of the organization we observed in mouse SC parallel that in mouse V1. In both areas, neurons are arranged in precise retinotopic maps and are robustly tuned to orientation and direction, yet nearby neurons do not form classical columns dominated by a single preference (Bonin et al., 2011; Chen et al., 2021; Inayat et al., 2015; Kaschube, 2014; Niell and Stryker, 2008). Instead, preferences are locally mixed, with only weak clustering at short distances. In V1, such clustering occurs on the scale of tens to hundreds of micrometers for orientation and, more modestly, for direction (Kondo et al., 2016; Ringach et al., 2016; Yu et al., 2025), comparable to the very small-scale clustering we found in SC. Local orientation biases in V1 have been linked to ON/OFF RF tiling in the retina (Jimenez et al., 2018), whereas SC may achieve better local coverage by receiving direction tuning from DSGCs and orientation tuning from OSGCs, or from pooling oppositely tuned DSGCs.

Theoretical accounts of map formation suggest that the absence of single-preference columns is expected in mouse SC. Across species, rodents and rabbits exhibit locally mixed tuning, whereas cats, ferrets, and primates display columnar maps (Kaschube, 2014), though not for all visual features (Martin and Schröder, 2013; Yen et al., 2007). Models based on the ratio of retinal inputs to cortical area—the mapping or sampling ratio—predict that low ratios favor a locally mixed organization, whereas high ratios favor columnar maps (Ibbotson and Jung, 2020; Jang et al., 2020; Najafian et al., 2022; Schmidt and Wolf, 2021). Published estimates place SC in a low-mapping-ratio regime: a larger fraction of RGCs projects to SC than to the thalamocortical pathway (Ellis et al., 2016; Martin, 1986), SC RFs are smaller than those in V1 (Niell and Stryker, 2008; Wang et al., 2010), and SC has a smaller surface area (Edwards et al., 1986; Mazade and Alonso, 2017). The same anatomical constraints that argue against classical columnar maps in mouse V1 therefore also argue against single-preference maps in SC.

### Functional implications and future directions

Biases toward directions aligned with body and gravity axes likely facilitate decoding of self-motion and ethologically relevant motion (Sabbah et al., 2017; Tiriac et al., 2022), whereas local mixing allows SC to represent many motion directions or edges at each location and support diverse behaviors (Hoy and Farrow, 2025; Wheatcroft et al., 2022). The spherical organization of OSGCs has been proposed to make it easy for a downstream decoder to infer edge orientation from the relative activity of a few OSGC types (Laniado et al., 2025). In the binocular zone, the vertical cell type of the longitudinal system and the horizontal type of the latitudinal system provide the highest decoding efficiency (Laniado et al., 2025). These two classes were the most common among sSC neurons recorded with Neuropixels in the binocular zone (Figure 7). This convergence indicates that SC neurons in the binocular zone preferentially represent particularly informative retinal orientation channels, even though these preferences remain weakly clustered at the neuronal level.

Future work should extend these measurements to larger SC populations while preserving circuit integrity. Ideally, this would combine broad retinotopic coverage, including anterior/binocular and posterior/monocular SC, with single-cell resolution. Optical methods are attractive but usually require removal of overlying cortex, which risks altering cortical input or damaging sSC; deep imaging approaches that access SC without cortical removal would therefore be especially valuable. New experiments should also quantify variability across individuals, given the substantial inter-animal differences reported previously, and test how organization generalizes to richer stimulus conditions. Natural images and movies, motion-contrast stimuli, and size- or context-dependent motion could reveal structure not engaged by simple gratings and bars (DePiero et al., 2024). Finally, relating SC organization to behavior—for example, by testing whether directions and orientations near overrepresented preferences are discriminated more accurately, or whether mice move head or eyes to bring stimuli into specific visual-field regions—will be useful to link the topographic organization in retina and SC to behavioral performance.

## Acknowledgements

We thank Profs Matteo Carandini (MC) and Kenneth D. Harris (KDH) for support of this project in their laboratory during data acquisition; we thank Charu Reddy and Charlotte Donald Wilson for help with mouse husbandry; we thank Michael Krumin for help with performing two-photon imaging experiments and Liad J. Baruchin for help with performing electrophysiological recordings; we thank Shai Sabbah for information provided regarding his geometrical model of retinal direction and orientation tuning; we thank Dr Luigi Federico Rossi and Prof Matteo Carandini for valuable feedback on the manuscript.

This work was supported by the People Programme (Marie Curie Actions) of the European Union’s Seventh Framework Programme (FP7/2007-2013) under REA Grant Agreement No 623872 (to SS), by the Biotechnology and Biological Sciences Research Council (BB/P003273/1 to MC and SS, BB/W014955/1 to SS), by Wellcome Trust grants (095669 and 205093 to MC and KDH), by a Sir Henry Dale Fellowship jointly funded by the Wellcome Trust and the Royal Society (220169/Z/20/Z to SS), by a Newton International Fellowship funded by the Royal Society (NIF\R1\211581 to MFGF), and by the Leverhulme Trust (doctoral fellowship to RG). For the purpose of open access, the author has applied a CC BY public copyright licence to any Author Accepted Manuscript version arising from this submission.

## Author contributions

Conceptualization and design: S.S.; data collection – two-photon imaging: S.S.; data collection and pre-processing – electrophysiology: M.F.G.F., R.G.; data analysis: S.S. and Z.H.; writing: S.S.; review and editing of manuscript: Z.H., M.F.G.F., R.G.

## Data and code availability

The pre-processed data used in this study are available at https://doi.org/10.6084/m9.figshare.30927041; code used to analyze pre-processed data is available at https://github.com/Schroeder-Lab/he-et-al_2025_tuning-topography. The raw data are available on reasonable request.

## Methods

### Experimental Model and Subject Details

All procedures were conducted in accordance with the UK Animals Scientific Procedures Act (1986) under personal and project licenses released by the Home Office following appropriate ethics review.

For two-photon imaging, we used 1 inbred C57Bl/6J (www.jax.org/strain/000664; 1 male) and 12 mice (7 females, 5 males) obtained by crossing Gad2-IRES-Cre (www.jax.org/strain/010802) and Ai9 (www.jax.org/strain/007909). The heterozygous offspring expressed TdTomato in glutamate decarboxylase 2-positive (GAD2+) cells to identify inhibitory neurons. For electrophysiological recordings, we used 4 Vgat-IRES-Cre mice (www.jax.org/strain/016962; 2 females, 2 males) and mice (1 female, 2 males) obtained by crossing Vgat-IRES-Cre (www.jax.org/strain/016962) and Ai14 (www.jax.org/strain/007908). Animals were 9–29 weeks old at the time of surgery and were used for experiments up to the age of 54 weeks. Mice were kept on a 12-h light: 12-h dark cycle. Most animals were single housed after the first surgery.

### Overview of collected data

|  | Retinal boutons (2P) | SC neurons (2P) | SC neurons (ephys) |
| --- | --- | --- | --- |
| Animals | 8 | 5 | 7 |
| Recording sessions | 17 | 12 | 23 |
| Recorded units | 3,677 | 4,769 | 1,474 |
| Units responsive to drifting gratings | 2,391 | 2,378 | 1,120 |
| Selective units | 603 DS-only;<br>232 OS-only;<br>643 DS & OS | 315 DS-only;<br>166 OS-only;<br>1,022 DS & OS | 124 DS-only;<br>358 OS-only;<br>165 DS & OS |
| Units for which RFs could be mapped | 1,979 of 3,282 recorded units (13 sessions, 8 animals) | 2,471 of 4,769 recorded units (12 sessions, 5 animals) | 539 of 1,120 recorded units (23 sessions, 7 animals) |
| Units with RF and tuned to direction or orientation | 674 (of 1,009 DS);<br>591 (of 769 OS) | 813 (of 1,337 DS);<br>697 (of 1,188 OS) | 172 (of 289 DS);<br>244 (of 523 OS) |

### Surgical procedures

Surgical procedures for animals undergoing two-photon imaging were described previously (Schröder et al., 2020). Here, we provide a shorter description and point out differences to animals undergoing electrophysiological experiments. Animals were anesthetized with isoflurane and the following drugs were administered subcutaneously: carprofen (5 mg/kg; Rimadyl, Pfizer) and dexamethasone (0.5 mg/kg; Colvasone, Norbrook) in animals undergoing surgery for two-photon imaging experiments (2P); or meloxicam (5 mg/kg; Metacam, Boehringer Ingelheim), buprenorphine (0.05 mg/kg; Vetergesic, Ceva Animal Health), and dexamethasone (4 mg/kg; Rapidexon, Dechra) in animals undergoing surgery for electrophysiology experiments (ephys). The scalp was shaved and disinfected, and local analgesia (2P: lidocaine: 6 mg/kg, Hameln pharmaceuticals ltd; ephys: bupivacaine: 7 mg/kg, Marcain, AstraZeneca) was injected subcutaneously under the scalp prior to the incision. Eyes were covered with eye-protective gel (Chloramphenicol, Martindale Pharmaceuticals Ltd). A local anesthetic (2P: 5% Lidocaine ointment, TEVA UK; ephys: 5% EMLA cream, Aspen) was applied to the ear bars, before the animal was placed into a stereotaxic apparatus. The skin covering and surrounding the area of interest was removed, and the skull was cleaned of connective tissue. A custom made headplate was positioned above the area of interest and attached to the bone with Superbond C&B (Sun Medical). Throughout all surgical procedures, the animal was kept on a heating pad to stabilize body temperature at 37°C. Subcutaneous injections of 0.01 ml/g/h of Sodium Lactate (Hartmann’s solution) were given. After the surgery, the animal was placed into a cage on a heating pad (37°C) for recovery from anesthesia. Mice were given three days to recover while being treated with Carprofen or Metacam.

To image retinal boutons in SC, we cloned a variant of GCaMP6f fused with a localization signal targeting synaptic terminals (SyGCaMP6f) and packaged it into an AAV2/2 vector (Addgene, 51085) so that GCaMP expression was restricted to boutons (Dreosti et al., 2009). We injected approximately 3 µl of AAV2/2.hSyn1.SyGCaMP6f.SV40 with a concentration of 2.44 × 10^12^ viral particles/ml into the vitreous humor of the left eye. In all animals used for two-photon imaging, a circular 4 mm craniotomy (centered at approximately −4.2 mm AP and 0.5 mm ML from Bregma) was made. For two-photon imaging of activity in SC neurons, we injected the virus AAV2/1.Syn.GCaMP6f.WPRE.SV40 (Chen et al., 2013) at a final concentration of 2.30–4.39e12 GC/ml, at a rate of 2.3 nl every 6 s (Nanoject II, Drummond), below the surface of the right SC. As the posterior SC is covered by a large sinus running through the dura, we implanted a custom-made implant permanently to push the sinus anteriorly and gain optical access to the SC. The implant was fixed to the skull with Vetbond (3M) and Superbond C&B (Sun Medical).

For electrophysiological recordings, a craniotomy (centered at −3.7 mm AP and 0.6 mm ML relative to Bregma, in the right hemisphere) was performed above the anterior SC at least one day before the first acute recording was performed.

### Two-photon imaging

Most of the two-photon imaging datasets have been used in a previous study (Schröder et al., 2020), and additional datasets have been collected in the same way. Shortly, two-photon imaging was performed using a standard resonant microscope (B-Scope, ThorLabs Inc.) equipped with a 16x, 0.8 NA water immersion objective (N16XLWD-PF, Nikon) and controlled by ScanImage 4.2 (Pologruto et al., 2003). Excitation light at 970–980 nm was delivered by a femtosecond laser (Chameleon Ultra II, Coherent). Multi-plane imaging was performed using a piezo focusing device (P-725.4CA PIFOC, Physik Instrumente, 400 µm range). Laser power was depth-adjusted and synchronized with piezo position using an electro-optical modulator (M350-80LA, Conoptics Inc.). Emission light was collected using two separate channels, one for green fluorescence (525/50 nm emission filter) capturing the calcium transients and one for red fluorescence (607/70 nm emission filter).

For imaging neurons in SC, we used 3–4 imaging planes separated by 9–30 µm at depths of 15–100 µm from the surface of SC. The field of view spanned 340–640 µm in both directions at a resolution of 512 x 512 pixels. The frame rate per plane was 6.0–7.5 Hz.

For imaging retinal boutons in SC we used 5 or 10 imaging planes (with 3 or 6 fly-back planes) with an inter-plane distance of <2 µm at depths of 8–46 µm from the surface of SC. The field of view spanned 42–135 µm and each plane was imaged at a rate of 7.5 Hz. Because the calcium indicator was localized to synaptic boutons, fluorescence was not affected by fibers of passage.

### Electrophysiology

Recordings were made using Neuropixels electrode arrays (Jun et al., 2017). Probes were mounted to a custom 3D-printed holder attached to a steel rod and positioned with a micromanipulator (uMP-4, Sensapex Inc.). For subsequent track reconstruction, probes were coated with a dye (DiI: ThermoFisher Vybrant V22888/V22885; or DiO: ThermoFisher Vybrant V22886) by immersing the shank for 5–10 min in an Eppendorf tube containing the dye solution, after which they were removed and air-dried. Probes had a soldered connection between external reference and ground; at the headstage, ground was connected to an Ag/AgCl wire placed on the skull. Craniotomies and the ground wire were kept covered with saline throughout the experiment. Probes were advanced through the dura and lowered to the target depth at ∼10 µm/s, then allowed to settle for 10 min before recording. Signals were acquired with SpikeGLX (https://billkarsh.github.io/SpikeGLX), and were separated into an action-potential (AP) band (hardware high-pass filtered at 300 Hz, sampled at 30 kHz) and a local-field-potential (LFP) band (low-pass filtered at 500 Hz, sampled at 2.5 kHz). Recordings were repeated across different penetration sites on multiple subsequent days.

### Experimental setup and visual stimuli

The mouse was head fixed with a headplate holder that did not obstruct the visual field. For two-photon imaging, the mouse was free to run on an air-suspended Styrofoam ball (20 cm in diameter), whose rotation was measured by two optical computer mice (Dombeck et al., 2010). For electrophysiology, the mouse was free to run on a plastic wheel (8.5 cm wide, 19 cm diameter) (Warren et al., 2021), whose rotation was measured by a rotary encoder (1,024 pulses per rotation, Kübler, Germany). The mouse was acclimated to head-fixation for at least three days before the first recording session, step-wise increasing fixation time from ≥5 min on the first day to ≥20 min on the last day.

For two-photon imaging, the mouse was surrounded by three computer screens (Iiyama ProLite E1980SD placed ∼20 cm from the mouse’s eyes; 60 Hz refresh rate) at right angles covering the visual field from −135° to 135° azimuth and −42° (bottom) to 42° (top) elevation. In some experiments, Fresnel lenses (BHPA220-2-6 or BHPA220-2-5, Wuxi Bohai Optics) were mounted in front of the monitors to compensate for reduction in luminance and contrast at steeper viewing angles relative to the monitors, and lenses were coated with scattering window film (frostbite, The Window Film Company) to prevent specular reflections. The red channel of the monitors was switched off to reduce light contamination in the red fluorescence channel. To track the eye contralateral to the recording site, we illuminated the eye with an infrared LED (850 nm, Mightex SLS-0208-A or Mightex SLS-0208-B). Videos of the eye were captured at 30 Hz with a camera (DMK 23U618 or DMK 21BU04.H, The Imaging Source) equipped with a zoom lens (Thorlabs MVL7000) and a filter (long-pass, Thorlabs FEL0750; or band-pass, combining long-pass 092/52×0.75, The Imaging Source, and short-pass FES0900, Thorlabs).

For electrophysiological recordings, visual stimuli were presented on one computer screen (Dell P1917S; 37.5 × 30 cm, 1280 × 1024 pixels, 60 Hz refresh rate), positioned 20 cm from the mouse’s left eye and spanning −95° (left) to 0° (right) azimuth and −27° (bottom) to 51° (top) elevation.

Brightness levels of the monitors were linearized around a mean level of 83 cd/m^2^ (two-photon imaging) or 36 cd/m^2^ (electrophysiology). All visual stimuli were defined in an orthonormal space, where X and Y represent azimuth (or longitude) and elevation (or latitude), respectively, in degrees of the visual field. To correctly display these stimuli, the 2D orthonormal space was mapped on the 2D planes of the monitors using the equirectangular or cylindrical mapping, which preserves the meridian lines as vertical lines, but introduces distortion around the poles. In the imaging experiments, the stimuli were created using PsychToolbox (Brainard, 1997) and then mapped onto screen using custom-written code. In the electrophysiology experiments, stimuli together with the correct mapping were produced using BonVision (Lopes et al., 2021).

Sinusoidal drifting gratings covered the complete screen space (full-field), had a spatial frequency of 0.08 cycles/deg, a temporal frequency of 2 cycles/s, and were presented at 100% contrast. Gratings drifted in one of 12 equidistant directions and were presented for 2 s separated by a gray screen for 3–6 s (two-photon imaging) or 2–3 s (electrophysiology).

Static sinusoidal gratings also covered the complete screen space, had a spatial frequency of 0.08 cycles/deg, and were presented at 100% contrast. Static gratings varied in orientation (0°–135°, in steps of 45°) and spatial phase (0°, 120°, 240°), comprising all possible combinations. Each static grating was presented for 1 s with an inter-stimulus interval of 2–3 s.

Moving bars were black on mean-gray background, had a width of 2.5°, a length spanning the full screen size, and moved at a speed of 15°/s. Bars varied in direction of movement (0°–315°, in steps of 45°). Each bar was presented in a window of 4 s with an inter-stimulus interval 3–5.5 s.

Within blocks of the same stimulus type, each stimulus was repeated 15 or 20 times following a pseudo-random presentation order.

To map RFs and to determine the position of SC along the probe, we presented checkerboard images (visual noise) with white, black and mean gray squares with an edge length of 10°. The stimulus updated at a rate of 6 Hz and was presented for >9 min. In each noise image, each square was randomly assigned its luminance value with a 98% probability of being gray and 1% probabilities of being black and white (for two-photon imaging), or 20 out of 288 squares was black or white (for electrophysiology). Alternatively, RFs were mapped using black and white circles with diameters varying between 0.5° and 32°. Circles were presented individually on gray background at a rate of 10 Hz, for at least 5 min.

### Data analysis

#### Preprocessing of two-photon imaging data

Preprocessing has been described previously (Schröder et al., 2020). Shortly, all raw two-photon imaging movies were analyzed using Suite2p (implemented in Matlab, Mathworks) to align frames and detect regions of interest (Pachitariu et al., 2016). We used the red channel reflecting TdTomato expressed in inhibitory neurons to align frames. For imaging data of retinal boutons, we also aligned frames in depth, i.e. when the brain moved perpendicular to the imaging planes, fluorescence data from neighboring imaging planes was used to correct this movement. Regions of interest (ROIs) were detected using the aligned frames of the green channel, and were then manually curated using the Suite2p GUI.

Using the aligned movies and detected ROIs resulting from Suite2p analysis, we extracted the fluorescence from the green and the red channel within each ROI. To correct the calcium traces for contamination by surrounding neuropil, we also extracted the fluorescence of the surrounding neuropil for each ROI using the green channel. To correct for contamination, the neuropil trace, N, was subtracted from the calcium trace, F, using a correction factor α: F_c_(t) = F(t) – α·N(t). The correction factor was determined for each ROI individually (Dipoppa et al., 2018). To correct for potential brain movements, we used the red fluorescence traces of each ROI to regress out changes in fluorescence that were not due to neural activity.

To avoid sampling the same neuron/bouton more than once, we removed duplicate ROIs that were close to each other in neighboring imaging planes and that had highly correlated calcium traces.

#### Spike sorting

Preprocessing and spike sorting were performed using the IBL-sorter pipeline (International Brain Laboratory et al., 2022), which includes common-median referencing, channel-noise whitening, detection and removal of noisy channels, and an initial estimate of probe drift. Kilosort2.5 was then executed using the detection thresholds of 9 for high-threshold detection and 3 for low-threshold detection. Unit quality was assessed with Bombcell (Fabre et al., 2023). Units were included only if they were labelled *good* by Bombcell and satisfied the following quality criteria: refractory-period violations ≤15%, <35% of spikes falling below the detection threshold, presence ratio >25%, and ≥300 spikes. All retained units were subsequently curated manually in phy (https://github.com/cortex-lab/phy) to identify potential merges or splits. Decisions were based on waveform similarity, the bimodality of waveform feature distributions, the spike amplitude distributions and drift patterns, and cross-correlogram structure between putative units.

#### Tracking of pupil position

Pupil videos were processed offline using custom MATLAB code as described previously (Schröder et al., 2020). Briefly, frames were mildly spatially filtered, the pupil contour was detected with a level-crossing edge detector, and artifacts from infrared reflections or whiskers were removed by excluding concave contour segments. Pupil position was then estimated from an ellipse fitted to the remaining contour. Eye blinks were detected from frame intensity and correlation to the average frame, visually checked, and excluded from further analysis.

#### Position of SC along recording probe

To identify the SC surface and the border between stratum griseum superficiale (SGS) and stratum opticum (SO) in Neuropixels recordings, we used visually evoked local field potentials (LFPs) following established approaches (Ito et al., 2017; Lee et al., 2020; Stitt et al., 2013; Zhao et al., 2014). For each recording, we first subtracted the median LFP across time from each channel to remove channel-specific offsets. We then processed the two vertical columns of Neuropixels channels separately. Within each column, we smoothed the raw LFP in time and depth by convolving with a Gaussian kernel (5 channels in depth, 21 time bins). At each time point, we subtracted the LFP averaged across all channels of a column to emphasize local depth-dependent structure. We next identified the most effective pixel (for noise stimuli) or circle (for circle stimuli) that resulted in the largest LFP amplitude and aligned the LFP to the pixel’s or circle’s onset times.

The SC surface was defined from the depth profile of the evoked LFP. For each channel depth, we averaged the aligned LFP across repetitions of the effective stimulus and then averaged pairs of channels at the same depth from the two columns, yielding one LFP trace per depth. For each depth, we computed the peak amplitude of the visually evoked LFP. Moving from the channel with the largest amplitude to superficial channels, the surface was assigned to the first channel whose evoked LFP amplitude fell below 25% of the peak amplitude.

To locate the SGS-SO border, we computed the current-source density (CSD) as the second spatial derivative of the stimulus-aligned LFP traces and averaged those across repetitions. At the latency with maximal positive CSD amplitude, we identified the first channel below this maximum whose CSD value was negative; this depth was taken as the SGS-SO boundary.

Because SO is part of superficial sSC, the SGS-SO border does not mark the boundary between sSC and dSC. We therefore used histological brain sections containing the probe tracks to estimate the depth of the sSC-dSC border. Probe tracks were registered to the mouse brain atlas (Franklin and Paxinos, 2007), and anatomical distances were measured along the reconstructed tracks. Across atlas-registered probe tracks, the full depth of sSC was 1.66 times the distance from the SC surface to the SGS-SO border. The total depth of SC was approximately 1.24 mm in the atlas. In addition, distances measured along the physical probes were 0.79 times the corresponding distances measured in the atlas. Therefore, for each recording, we defined the SC surface and SGS-SO border from the visually evoked LFP and CSD as described above, defined the sSC-dSC border as 1.66 times the surface-to-SGS-SO distance, and defined the bottom of SC as 1.24 mm × 0.79 below the SC surface.

#### Response amplitudes to visual stimuli

The calculation of response amplitudes in response to drifting gratings has been described previously (Schröder et al., 2020). Here, we used the same strategy for responses to static gratings and moving bars. We quantified response amplitudes to the stimuli by modelling the calcium trace as the sum of stimulus-evoked responses. For each stimulus presentation, the response was modeled as the product of a scaling factor and a temporal response kernel that was 10 s long and aligned to stimulus onset. The same temporal kernel was used for each trial independent of specific stimulus features such as direction of movement. Compared with simply subtracting the mean pre-stimulus activity from the mean activity during stimulus presentation, this approach models all stimulus presentations in one continuous trace, allowing residual calcium signals from preceding trials to be separated from the response evoked by the current stimulus.

The model was fitted iteratively. In the first step, we fitted the temporal kernel using a General Linear Model, i.e. a linear regression model of the form

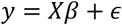

where *y* is the fluorescence trace of one ROI, *X* is the design matrix, *β* contains the fitted temporal kernel, and *ε* is the residual error. *X* has size *T* × *K*, where *T* is the number of samples in each calcium trace and *K* is the number of samples in each kernel. *X_i_*_,*j*_ equals the current response amplitude *a_r_* (initially set to 1) for trial *r* if trial *r* started *i* − *j* samples before time point *i*, and otherwise *X_i_*_,*j*_ = 0. The resulting kernel *β* was normalized to have an absolute maximum of 1.

In the second step, we fitted the response amplitudes in a second General Linear Model

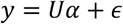

where *y* is again the fluorescence trace of one ROI, *U* is the design matrix for fitting trial amplitudes, *α* contains the fitted trial amplitudes, and *ε* is the residual error. If T is the number of samples in the calcium trace and R is the number of grating presentations, then *U* has size *T* × *R*, and *α* has size *R* × 1. Each column of *U* corresponds to one trial. Specifically, if trial *r* started at sample *s*, then

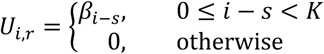

where *β* is the fitted temporal kernel of length *K*.

Both steps were repeated until the values of the response amplitudes stopped changing across iterations.

To estimate temporal kernels and amplitudes in response to moving bars, we introduced a third step in the iterative fitting procedure to determine the delay of the response relative to stimulus onset, which depended on motion directions and the location of the RF. The delays were estimated for each motion direction using the cross-correlation between the trial-averaged response and the trial-averaged prediction from the current response kernel and trial amplitudes.

To test responsiveness of a unit to a stimulus type, i.e., whether it had a stimulus response that was significantly different from baseline, we randomly shifted the response of each unit against the stimulus presentation times. A unit was determined to be *responsive* if the mean squared error (MSE) of the fits to the measured responses was smaller than the 95% confidence interval of MSEs resulting from fits to the randomly shifted responses.

To estimate response amplitudes from electrophysiological recordings, we calculated firing rates from spikes between 0 and 2 s after stimulus onset and, for each trial, subtracted the firing rate between 0 and 0.5 s before stimulus onset. The unit was deemed *responsive* to a stimulus type if the grand mean response was different from zero (using a linear model and a t-test on the intercept) or if responses to different stimuli of the same type were significantly different from each other (using a linear model and an F-test on the coefficients for all stimuli).

#### Direction and orientation preferences and selectivity index

The direction and orientation selectivity index of a unit was determined from its mean response amplitudes, *R_k_*, to each stimulus (drifting gratings, static gratings, moving bars). Mean amplitudes of units suppressed by gratings (responses to most stimuli <0) were inverted; remaining negative amplitudes were set to zero. Direction selectivity was assessed by scaling unit vectors pointing into the stimulus direction, *α_k_*, by the mean amplitudes, then summing these vectors: 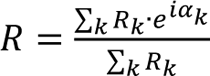. The angle and length of *R* are preferred direction and direction selectivity index (DSI). To determine preferred orientation and orientation selectivity, *α_k_*was first doubled. Significance of direction and orientation selectivity, respectively, was determined by a permutation test (1,000 permutations of stimulus direction or orientation across trials). Units were classified as direction-or orientation-selective only if the corresponding selectivity index was significantly larger than zero and at least 0.1. Colormaps for the visualization of preferred directions and orientations are perceptually uniform (Kovesi, 2015).

#### Mapping of receptive fields

We presented sparse sequences of white and black squares, or white and black circles of varying sizes, to characterize the receptive fields, using the responses to white squares or circles for the ON subfield and to black squares or circles for the OFF subfield. For imaging data, we first cleaned the calcium traces by removing any slow decays at the start of experiments (likely introduced due to bleaching) and high-pass filtering the data (subtracting smoothed traces treated with a moving median with a window size of 20 s). We used linear regression to simultaneously fit two spatio-temporal filters—one for the ON field and one for the OFF field. The time dimension of the filters spanned 0.2–0.5 s after each stimulus frame for two-photon data, and 0–0.2 s for electrophysiology data. For the ON field, only the appearance of white squares/circles was considered, for the OFF field only black squares/circles. The fits were regularized by imposing spatial smoothness on the two receptive fields (Smyth et al., 2003). We then fitted 2D Gaussians to the ON-, the OFF- and the mean between the ON- and OFF-subfield. The final spatial RF was the Gaussian that minimized the mean squared error with the mapped RF (if only one subfield was fitted by the Gaussian, the fit for the other subfield was set to zero for each pixel). The final temporal filter of the RF was determined by the least-squares scalar weights of the fitted 2D Gaussian for each time point.

A unit was considered to have a genuine receptive field if (1) the explained variance for the response to the visual noise predicted by the final spatio-temporal RF (Gaussian weighted in time) was ≥0.01, and (2) the peak of the fitted Gaussian was at least 5 times the SD of the residuals between mapped RF and fitted Gaussian.

#### Correction for eye movements during receptive field mapping

To assess whether small eye movements during head-fixed two-photon imaging affected the estimated RF positions, we re-fitted RFs while accounting for concurrent gaze position on each stimulus frame. We modelled the mapping from pupil displacement (in pixels, relative to the median pupil position) to gaze shift in visual degrees as a linear transform, [*Δaz*; *Δel*] = *A* · [*Δx*; *Δy*], where *A* is a 2 × 2 matrix estimated from the data. The elements of *A* were constrained to reflect the expected geometry of mouse eye movements: horizontal pupil displacements were required to produce larger shifts in azimuth than in elevation, vertical displacements to produce larger shifts in elevation than in azimuth, and small rotations between the camera and visual coordinate systems were permitted.

We estimated *A* and the gaze-corrected RF centers jointly using an iterative alternating procedure. In step 1, we shifted each frame of the sparse noise stimulus by the gaze displacement predicted under the current *A*, using bilinear interpolation of the stimulus grid, and re-fitted the spatio-temporal RF maps using the same regularized ridge regression followed by 2D Gaussian fitting, as described in Mapping of receptive fields. In step 2, holding the Gaussian RF shapes fixed, we re-optimized *A* by minimizing the sum of squared errors between the observed neural responses and those predicted by convolving each Gaussian RF with the original stimulus displaced by *A* · [*Δx*; *Δy*] on each frame, using constrained nonlinear optimization (MATLAB fmincon). The two steps were repeated with the updated *A* if doing so reduced prediction error; otherwise the algorithm was terminated. Iterations continued until the mean squared change in the elements of *A* fell below 10⁻⁵, or after a maximum of 10 iterations.

#### Relating RF locations to retinotopy

To relate the positions of imaged units within the FOV to their RF locations in visual space, we estimated a retinotopic transform for each recording. We modeled azimuth and elevation of the RF center as linear functions of the unit’s position in the imaging plane and fitted this model under the constraint that azimuth and elevation axes in brain space were orthogonal.

We only considered units with a genuine RF (see RF mapping) and a recording had to have at least 10 such units. Units with outlier RF locations (elevation or azimuth was >5 median absolute deviations away from the median across the population) were excluded. We then fitted a linear mapping from brain coordinates (*x*, *y*) to RF azimuth *az* and elevation *el*:

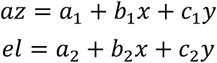

To enforce orthogonality between the azimuth and elevation axes in brain space, we constrained the elevation coefficients (*b*_2_, *c*_2_) to be orthogonal to the azimuth coefficients (*b*_1_, *c*_1_):

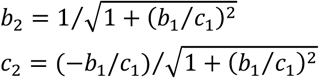

Model parameters were obtained by weighted least squares, weighting each unit by the variance explained by its final spatio-temporal RF model, so that units with more reliable RF estimates contributed more strongly to the fit. The resulting transform defined the optimal scaling, translation, and rotation that minimized the distance between mapped RF locations and the fitted retinotopic coordinates for each recording.

#### Geometric model of direction and orientation preferences

To relate measured preferences to the retinal topographic organization of directions and orientations, we used geometric models derived from retinal recordings (Laniado et al., 2025; Sabbah et al., 2017). For each unit, we computed four predicted directions and four predicted orientations at its RF location, corresponding to the four motion classes and four orientation classes defined in these studies.

Both retinal studies expressed optimal axes in retinal coordinates and then transformed them into body coordinates based on measurements of eye and head position. Axes for the left eye were provided (personal communication from Shai Sabbah) as points on the unit sphere in azimuth-elevation coordinates, where elevation is 0° at the horizon and increases towards the top of the visual field, and azimuth is 0° at the vertical meridian (nose) and increases clockwise. For motion directions, the four axes pointed to the convergence points of directional vectors on the sphere (advance, retreat, rise, fall): advance = (180°, −15°), retreat = (0°, 11°), rise = (−67°, −83°), fall = (106°, 88°). For orientations, we used four axes defining two longitudinal (“long”) and two latitudinal (“lat”) fields. Longitudinal orientations: horizontal = (187°, 90°), vertical = (98°, 26°); latitudinal orientations: horizontal = (74°, 22°), vertical = (232°, 85°).

These axes were defined under the assumption that the mouse head is pitched downward by 29° in the ambulatory posture (Oommen and Stahl, 2008), with 0° defined as bregma and lambda at the same height. In our experiments, mice were head-fixed and generally positioned with bregma lower than lambda to improve access to the SC, but the exact head pitch was not recorded for each mouse when the headplate was fixed during surgery. We therefore rotated all axes about the interaural (y) axis by an angle *α*, constrained to the range 0–29°, to account for uncertainty in head pitch. For each mouse, *α* was optimized by minimizing the discrepancy between predicted and measured direction and orientation preferences. The fitted *α* values for each mouse and recording modality are shown in Figure S3I. Rotation on the sphere proceeded as follows:

1. Each axis with azimuth *T_az_* and elevation *T_el_* was converted to Cartesian coordinates:

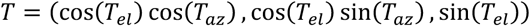
2. A rotation by angle *α* around the +y axis (right-hand rule) yielded:

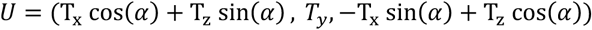
3. The result was then transformed back to azimuth-elevation coordinates:

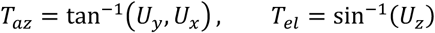

We obtained the predicted local direction or orientation at a given RF location as follows:

1. We converted the rotated axis (*T_az_*, *T_el_*) and the RF position (*r_az_*, *r_el_*) to Cartesian vectors:

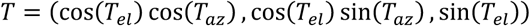

and

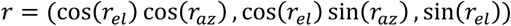
2. For a longitudinal field, the local direction vector *v* at *r* was the component of *T* tangential to the sphere:

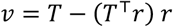
3. For a latitudinal field, *v* was given by the cross-product:

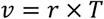
4. We then projected *v* onto the tangent plane at *r* using basis vectors aligned with increasing azimuth and elevation:

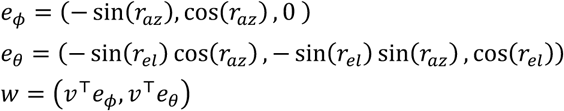
5. We computed the local direction angle *x* on the sphere:

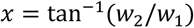
6. Finally, we converted *x* to our convention (0° = rightward, 90° = downward) as:

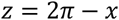

Using these constructions, we obtained, for each unit and each model (direction and orientation), the predicted vectors corresponding to the four direction classes and four orientation classes at the unit’s RF. For each unit, we then assigned the predicted direction and orientation whose vector was closest (smallest angular difference) to the measured preferred direction and orientation. This angular difference defined the prediction error between the unit’s preference and the geometric model. To test whether the retinal geometric model explained the data better than expected by chance, we compared the median angular difference across units to a null distribution obtained by permuting RF locations 1,000 times and report the median together with the 2.5th–97.5th percentile interval of this permutation distribution.

## Quantification and Statistical Analysis

All tests were two-sided, unless otherwise stated.

Permutation tests were performed by generating surrogate datasets from the measured data, computing the test statistic for each surrogate dataset, and comparing the resulting null distribution to the test statistic of the measured data, i.e. whether or not the measured test statistic falls inside the 2.5–97.5 percentiles of the null distribution. The surrogate datasets were generated by randomly permuting the relevant feature/label across sample points.

## Supplementary Figures

**Figure S1.**
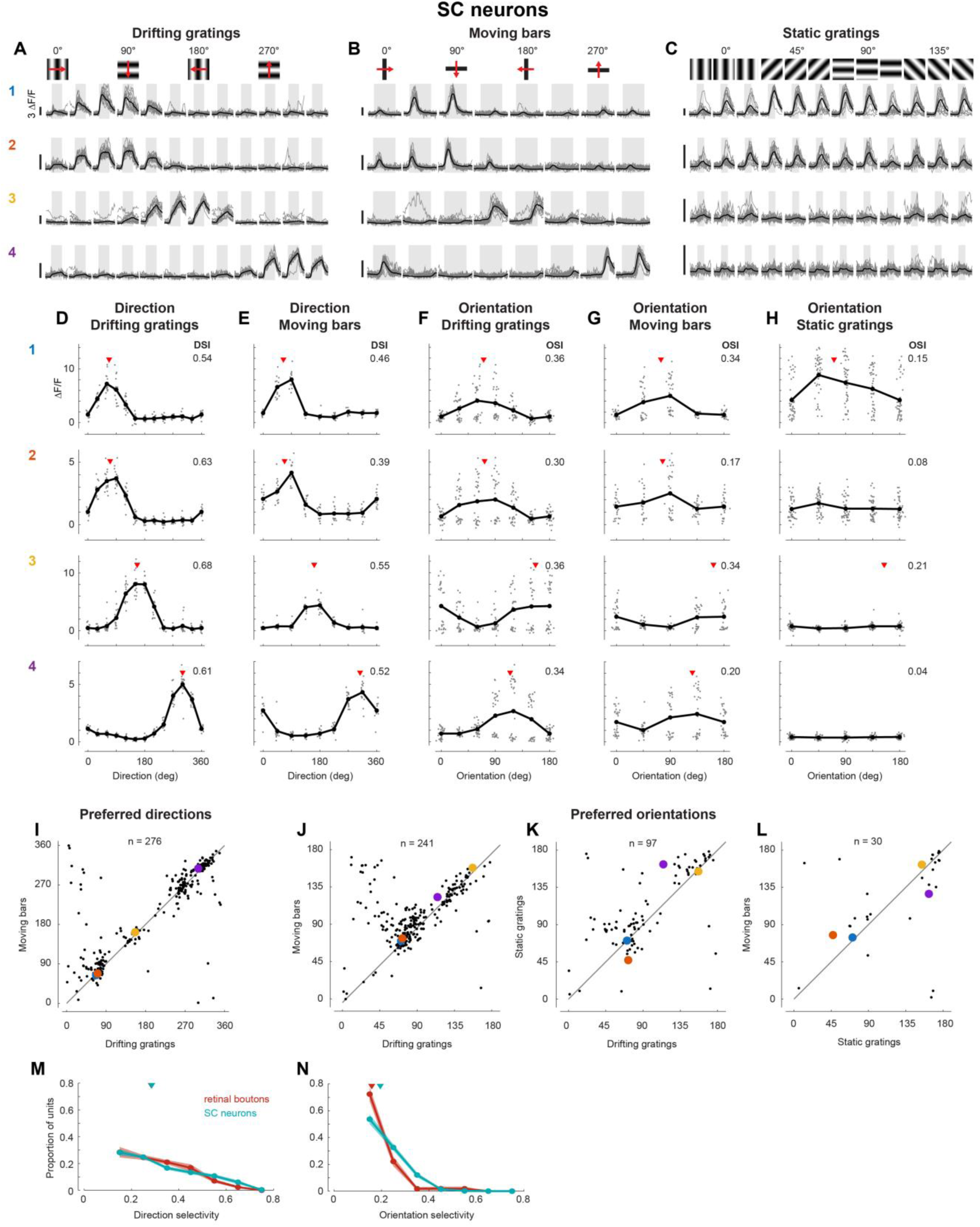
Direction and orientation tuning of SC neurons were consistent across stimulus types. A–C. Single-trial (gray) and mean fitted (black) calcium traces of four example neurons in response to drifting gratings (A; 12 directions, 2 s each), moving bars (B; 8 directions, 4 s each), and static gratings (C; 4 orientations × 3 spatial phases, 1 s each). Stimulus epochs are indicated by gray shading. We recorded 2,283 SC neurons in response to moving bars and 2,314 neurons in response to static gratings, both within 6 sessions across 5 mice. As for drifting gratings, the kernel fits matched the data very well (mean ± SD explained variance: 0.33 ± 0.21 for bars, 0.30 ± 0.20 for static gratings). Across SC neurons, responsiveness dropped from 50% for drifting gratings to 31% (697/2,283) for moving bars and 41% (945/2,314) for static gratings. D–H. Direction (D, E) and orientation (F–H) tuning curves for the example neurons in A–C, showing single-trial response amplitudes (gray dots), mean amplitudes (black dots), and preferred direction or orientation (red triangles) when tuning was significant. Tuning is shown for drifting gratings (D, F), moving bars (E, G), and static gratings (H). 62% (434/697) of neurons responsive to moving bars were selective for direction (compared to 56% for drifting gratings), while 54% (373/697) were selective for orientation (compared to 50% for drifting gratings). In contrast, only 24% (224/945) of neurons responding to static gratings were selective for orientation. I–L. Direct comparisons of preferred directions (I) and preferred orientations (J–L) obtained from different stimulus classes. Colored dots highlight the example neurons from A–C. SC neurons that were significantly tuned in response to several visual stimuli exhibited highly coherent preferences for direction and orientation with an average difference of 3.0° in preferred direction for drifting gratings and moving bars, and average differences of up to 10.0° in preferred orientation (drifting gratings vs bars: 3.8°; drifting vs static gratings: 10.0°; bars vs static gratings: 1.8°). M, N. Mean distribution (± SEM) of direction (M) and orientation (N) selectivity measured with drifting gratings, across all sessions with at least 20 selective units. Triangles mark the median (across sessions) of the median selectivity index.

**Figure S2.1.**
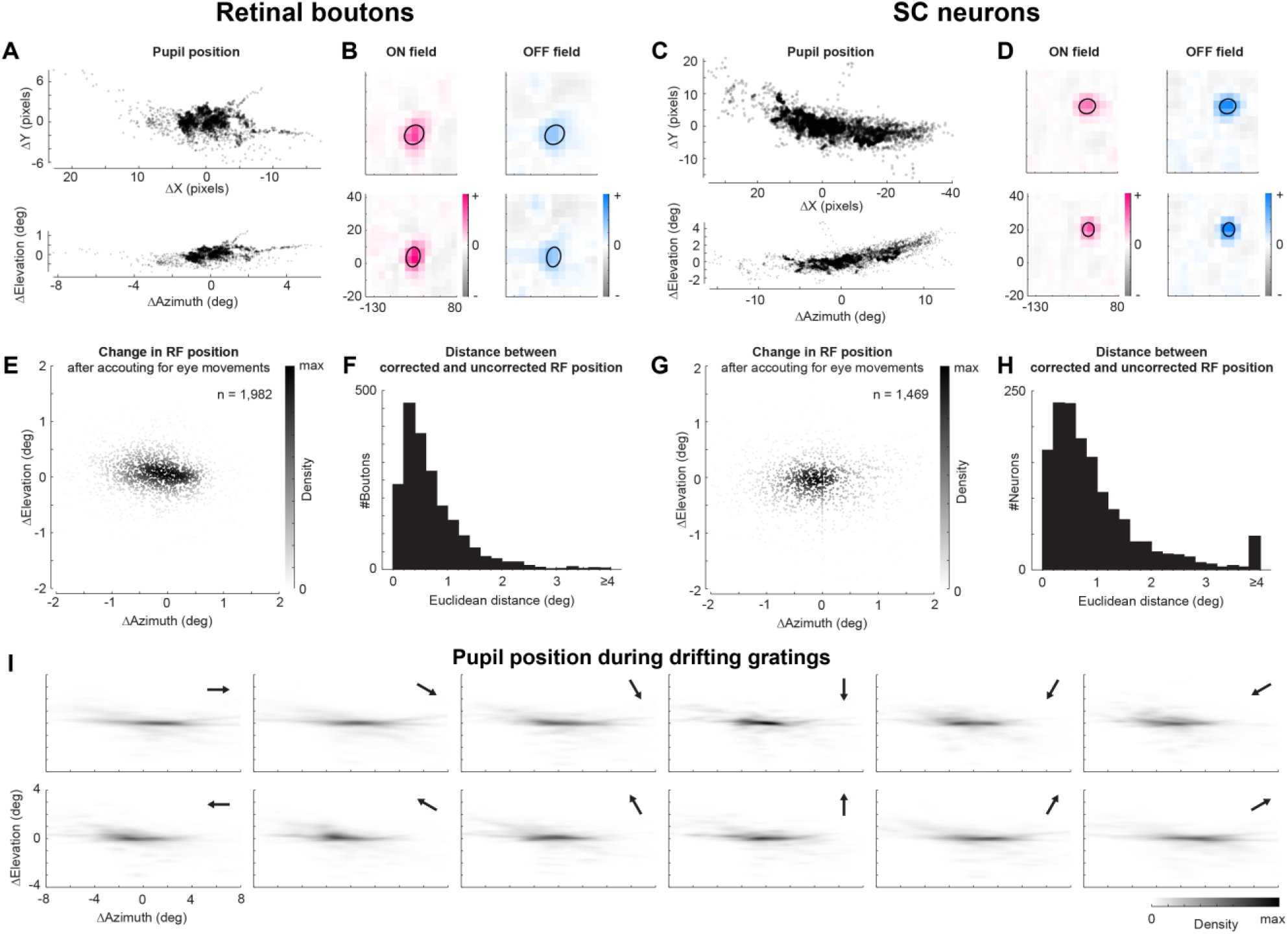
Eye movements did not substantially affect receptive field position estimates. A, C. Example distributions of pupil position during sparse noise stimulation, for a retinal bouton session (A) and an SC neuron session (C). Top: pupil displacement in camera pixels (ΔX, ΔY). Bottom: gaze displacement in visual degrees (ΔAzimuth, ΔElevation), obtained by applying the fitted linear transform A (see Methods). B, D. Example ON and OFF receptive fields fitted without (top) and with (bottom) eye-position correction, for the sessions shown in A and C. Ellipses indicate the fitted 2D Gaussian. Color scales are matched between corrected and uncorrected maps. E, G. Change in RF center position after eye-position correction (corrected minus uncorrected) for all well-fitted retinal boutons (E) and SC neurons (G). Point color indicates local density. F, H. Distribution of Euclidean distances between corrected and uncorrected RF centers for retinal boutons (F) and SC neurons (H). **I.** Gaze position distributions during drifting grating stimulation, shown separately for each of the 12 grating directions (arrows). Each panel shows a kernel density estimate pooled across all sessions for which pupil data were available (n = 21 sessions).

**Figure S2.2.**
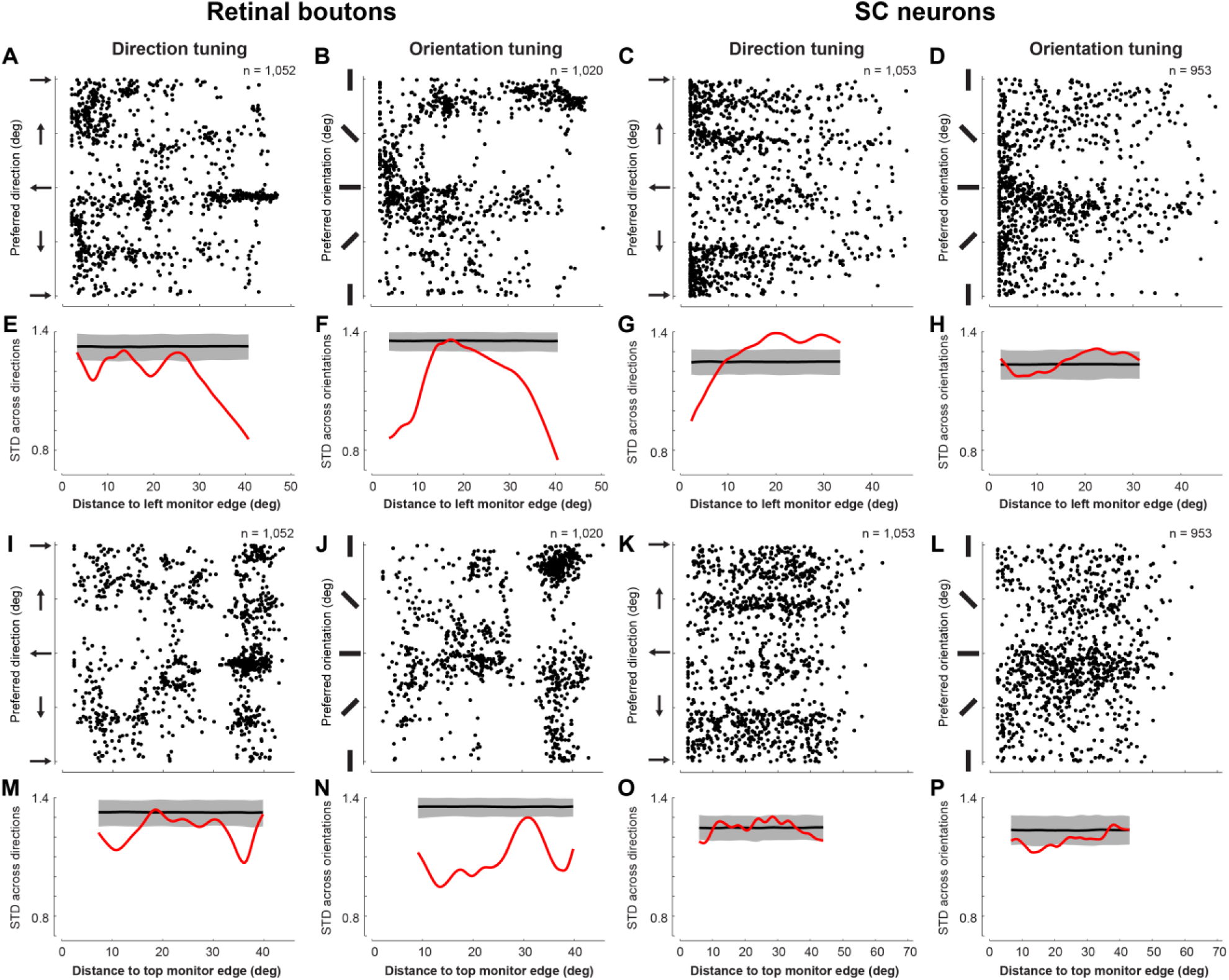
Proximity to monitor edges did not systematically affect measured tuning preferences. A–D. Direction (A) and orientation (B) preferences versus distance from the RF center to the nearest vertical (left/peripheral) monitor edge, for all selectively tuned retinal boutons with mapped RFs. Corresponding data for SC neurons in C and D. E–H. Circular SD (red) of preferred directions (E) and orientations (F) for retinal boutons as a function of RF distance to nearest vertical monitor edge, compared to a null distribution obtained by permuting edge distances (black line: median; gray band: 2.5th–97.5th percentile interval). Corresponding data for SC neurons in G and H. If the monitor edge had biased tuning preferences towards a specific direction or orientation, we would expect that preferences of units closer to the monitor edge are more similar to each other than of units farther away from the edge, which was not the case in our data. I–P. As in A–H, but using distance from the RF to the nearest horizontal (top) monitor edge.

**Figure S3.**
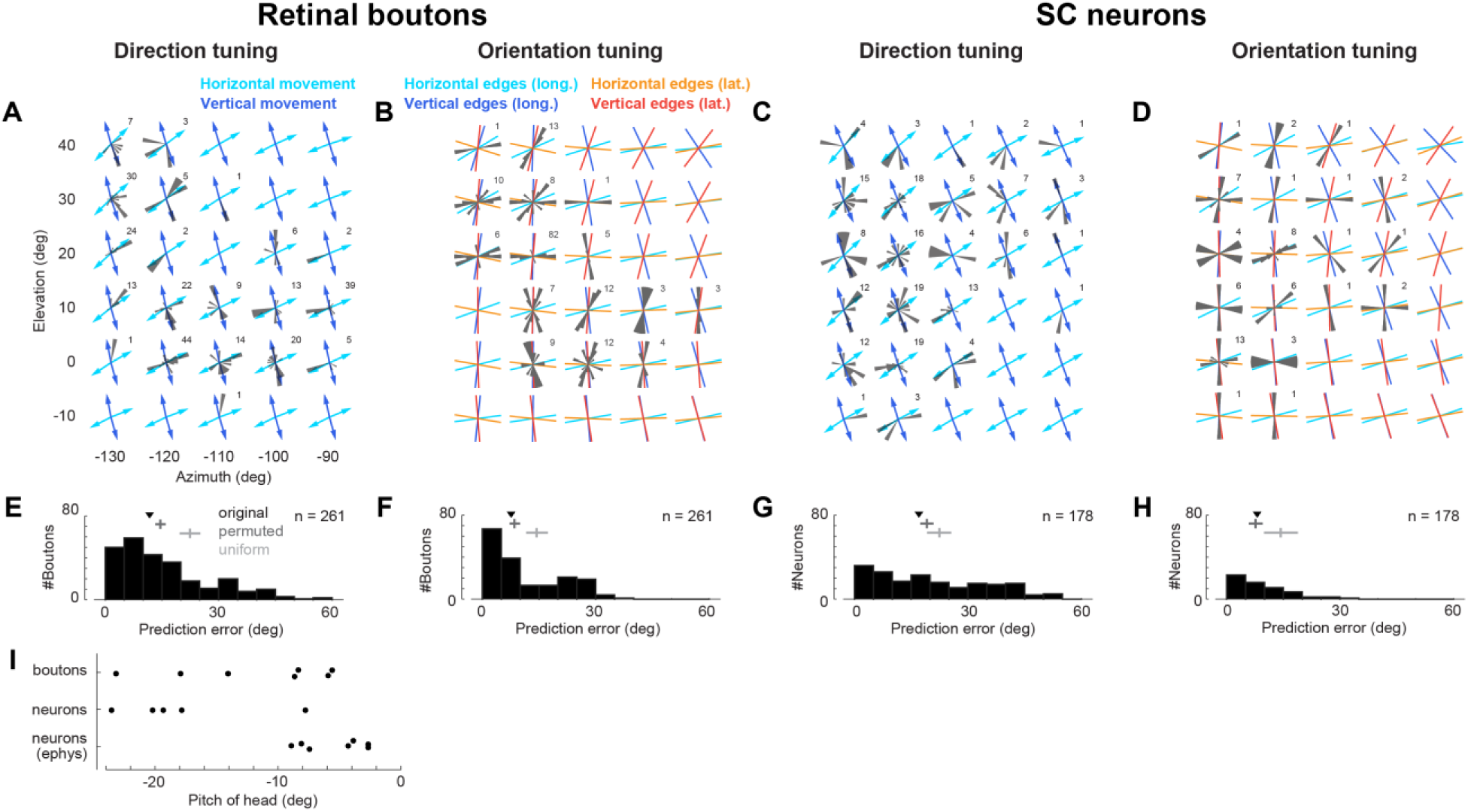
Topography for DS-only and OS-only units is similar to overall population. A–D. Direction (A) and orientation (B) preferences of DS-only and OS-only retinal boutons across visual space (RFs mapped, not retinotopic). Arrows in A indicate the predicted DSGC longitude directions; colored lines in B indicate predicted OSGC longitudes and latitudes (see Figure 3A–C). Numbers indicate population size in each polar histogram. Corresponding data for SC neurons in C and D. E–H. Angular differences between measured preferences and the closest predicted directions (E) or orientations (F) for DS-only and OS-only retinal boutons. Median differences (triangles) are compared to null distributions obtained by permuting RF locations (dark gray; median and 2.5–97.5 percentile intervals) or by drawing preferences from a uniform distribution (light gray; median and 2.5–97.5 percentile intervals). Direction: 11.7° original, 13.2°–16.0° permuted, 19.7°–25.2° uniform; orientation: 7.6° original, 7.2°–10.0° permuted, 11.6°–17.3° uniform. Corresponding data for SC neurons in K and L. Direction: 17.0° original, 17.4°–20.9° permuted, 19.2°–25.6° uniform; orientation: 8.1° original, 5.9°–9.6° permuted, 9.9°–18.9° uniform. Direction preferences of retinal boutons matched model predictions significantly better than those of SC neurons (p = 0.0016, Wilcoxon rank sum test). Orientation preferences matched model predictions to equal degrees in retinal boutons and SC neurons (p = 0.49, Wilcoxon rank sum test). **I.** Optimal head pitch angles estimated for each animal and recording modality. For each mouse, head pitch was chosen as the value that minimized the difference between measured direction/orientation preferences and the corresponding retinal topographic model predictions.

**Figure S4.**
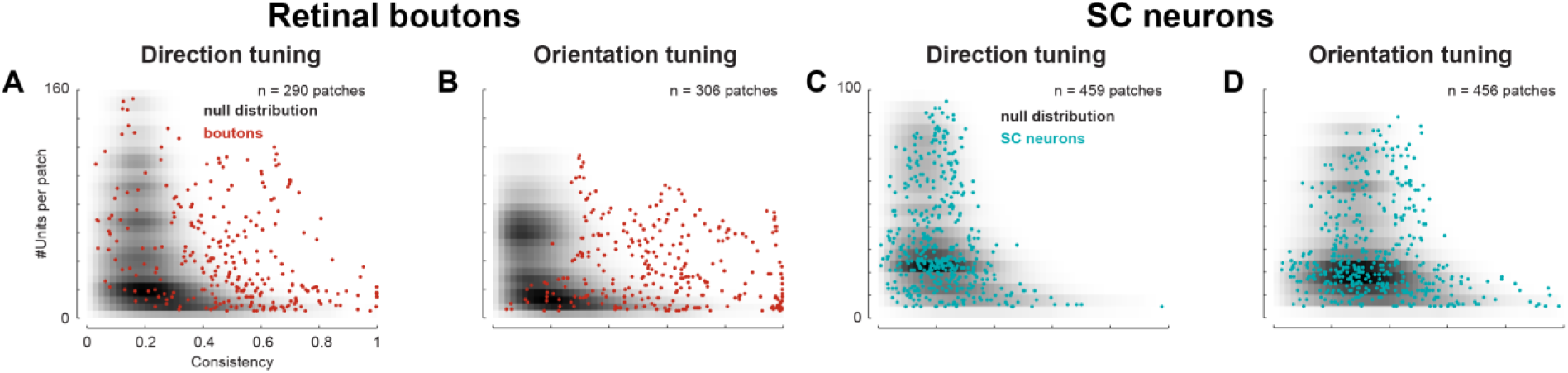
Local consistency is not trivially explained by the number of sampled units. A–D. Mean consistency of direction (A) and orientation (B) preferences within each 10° patch, plotted versus the number of retinal boutons in that patch (red dots). The background grayscale indicates the density of units from a null distribution obtained by permuting RF locations across boutons. Corresponding data for SC neurons in C and D.

**Figure S5.**
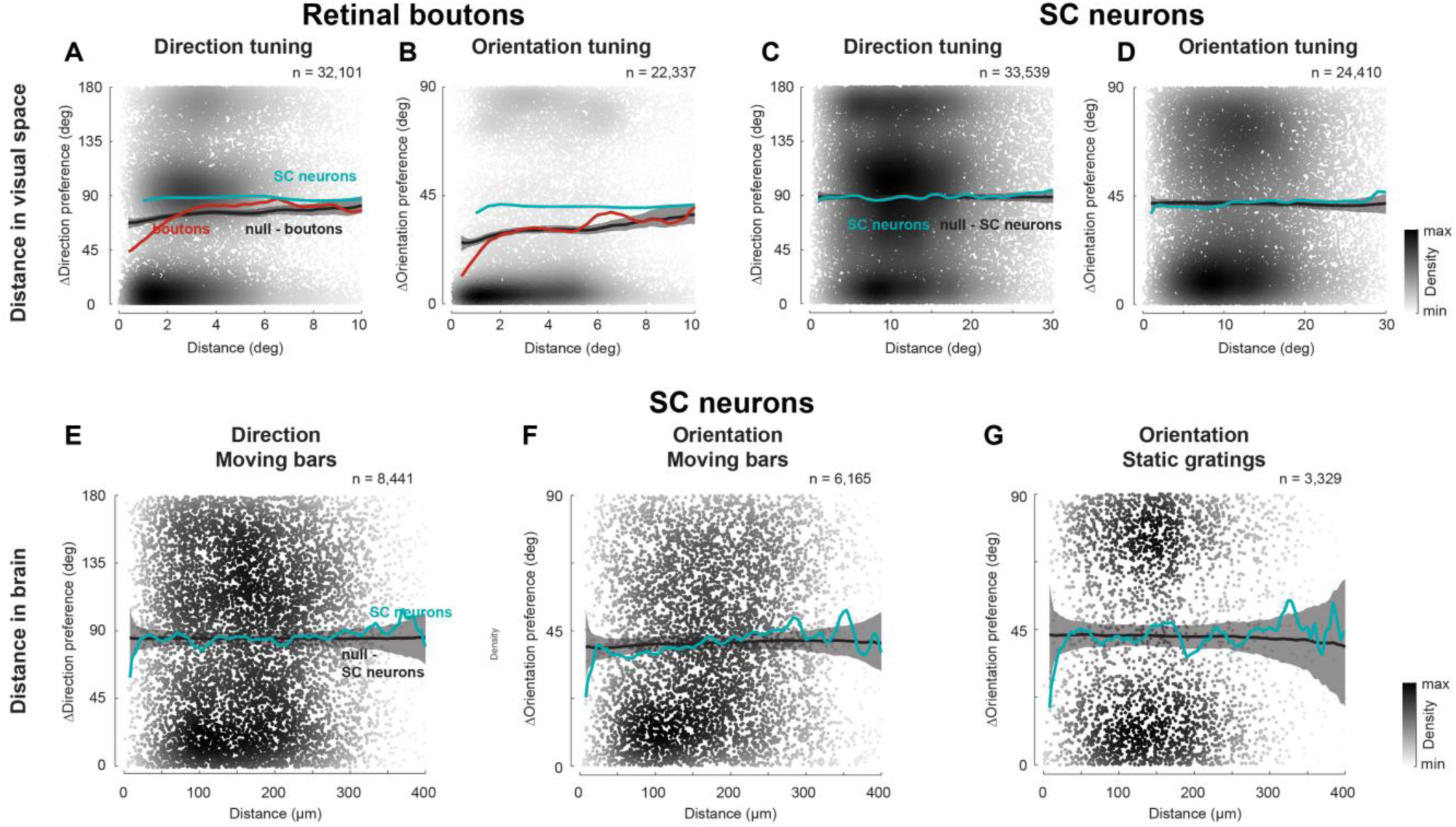
Weak clustering persisted with RF-based distances and under different stimulus conditions. A–D. Pairwise difference in direction (A) and orientation (B) preferences versus distance between mapped RFs for all pairs of simultaneously recorded retinal boutons. For comparison, mean pairwise differences for SC neurons (teal) are overlaid (same data as in C, D). Corresponding data for SC neurons in C and D. Only retinal boutons with RFs closer than 2.2 visual degrees had direction and orientation preferences that were more similar than expected from the null distribution. Tuning differences for SC neurons were much closer to the null distribution. Only orientation preferences were significantly smaller for neurons with RF distances of up to 10 visual degrees. E. Pairwise difference in direction preferences versus lateral distance in SC (depth ignored) for all pairs of simultaneously recorded SC neurons, with preferences measured from responses to moving bars. Mean pairwise tuning differences (86°) were close to those expected from uniform distributions. F, G. Same as in E, for orientation preferences measured from moving bars (F) and static gratings (G). Mean pairwise tuning differences were close to those expected from uniform distributions (40° for moving bars, 43° for static gratings). In all panels, dots are colored by density, and mean difference as a function of distance is compared to null distribution obtained by permuting neural or RF locations (black line: median; gray band: 2.5th–97.5th percentile interval).

**Figure S6.**
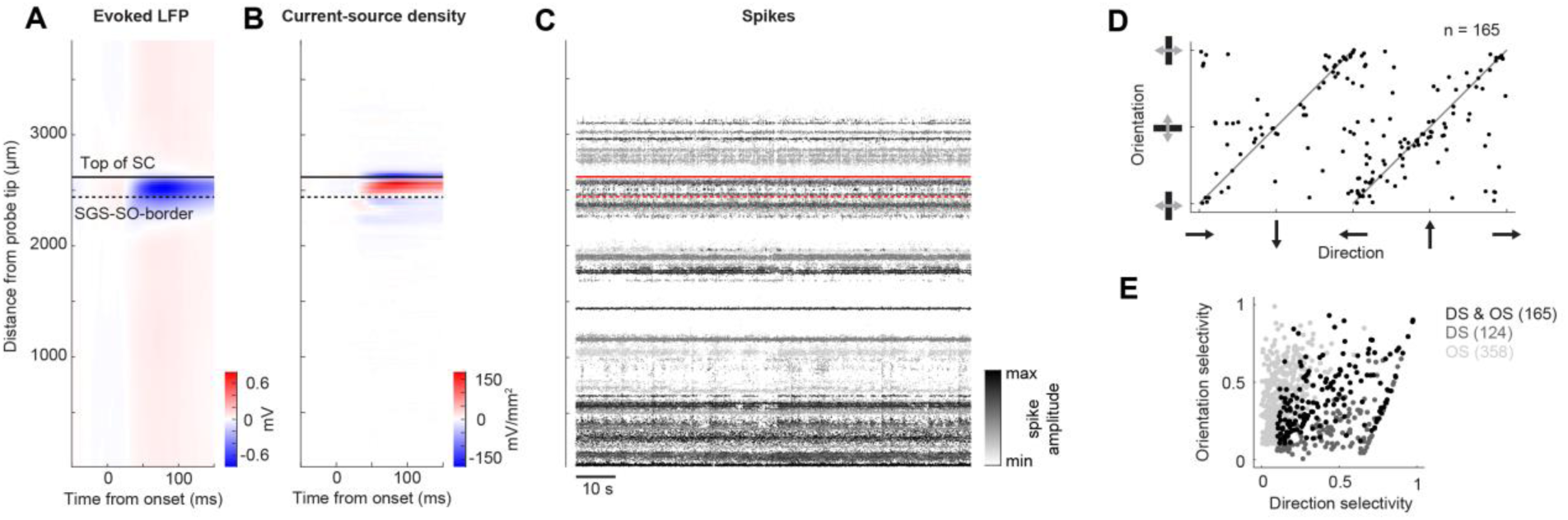
Determining position of SC along Neuropixels probe and tuning properties in electrophysiology data. A, B. Evoked local field potential (LFP, A) and current-source-density (CSD, B) profiles from an example recording in response to the most effective square of the visual noise stimulus. The surface of SC is defined at 25% of the peak amplitude in the evoked LFP. The border between SGS and SO is defined as the zero-crossing below the largest positive CSD peak. **C.** Spike amplitudes at spike times of all single units in the same recording as in A and B, plotted against their depths. **D.** Preferred direction versus orientation for SC units selective for both features. **E.** Direction versus orientation selectivity for all tuned SC units.

## References

Ahmadlou M, Heimel JA. 2015. Preference for concentric orientations in the mouse superior colliculus. Nature communications 6:6773. DOI: 10.1038/ncomms7773, PMID: 25832803

Baden T, Berens P, Franke K, Román Rosón, M, Bethge M, Euler T. 2016. The functional diversity of retinal ganglion cells in the mouse. Nature 529:345–350. DOI: 10.1038/nature16468

Basole A, White LE, Fitzpatrick D. 2003. Mapping multiple features in the population response of visual cortex. Nature 423:986–90. DOI: 10.1038/nature01721, PMID: 12827202

Bonin V, Histed MH, Yurgenson S, Reid RC. 2011. Local diversity and fine-scale organization of receptive fields in mouse visual cortex. The Journal of Neuroscience 31:18506–21. DOI: 10.1523/JNEUROSCI.2974-11.2011, PMID: 22171051

Borges K, Xian Q, Lu R, Liang Y, Haley S, Nigam S, Ji N. 2025. Retinal ganglion cell input to superior colliculus encodes salient information. DOI: 10.1101/2025.11.06.687035

Brainard DH. 1997. The Psychophysics Toolbox. Spatial Vision 10:433–436. PMID: 9176952

Cang J, Chen C, Li C, Liu Y. 2024. Genetically defined neuron types underlying visuomotor transformation in the superior colliculus. Nature Reviews Neuroscience 25:726–739. DOI: 10.1038/s41583-024-00856-4

Chen H, Savier EL, DePiero VJ, Cang J. 2021. Lack of Evidence for Stereotypical Direction Columns in the Mouse Superior Colliculus. The Journal of Neuroscience 41:461–473. DOI: 10.1523/JNEUROSCI.1155-20.2020

Chen T-W, Wardill TJ, Sun Y, Pulver SR, Renninger SL, Baohan A, Schreiter ER, Kerr RA, Orger MB, Jayaraman V, Looger LL, Svoboda K, Kim DS. 2013. Ultrasensitive fluorescent proteins for imaging neuronal activity. Nature 499:295–300. DOI: 10.1038/nature12354, PMID: 23868258

De Franceschi G, Solomon SG. 2018. Visual response properties of neurons in the superficial layers of superior colliculus of awake mouse. The Journal of Physiology. DOI: 10.1113/JP276964

De Malmazet D, Kühn NK, Farrow K. 2018. Retinotopic Separation of Nasal and Temporal Motion Selectivity in the Mouse Superior Colliculus. Current Biology 28:2961–2969.e4. DOI: 10.1016/j.cub.2018.07.001, PMID: 30174186

De Malmazet D, Kühn NK, Li C, Farrow K. 2024. Retinal origin of orientation but not direction selective maps in the superior colliculus. Current Biology 34:1222–1233.e7. DOI: 10.1016/j.cub.2024.02.001, PMID: 38417446

DePiero VJ, Deng Z, Chen C, Savier EL, Chen H, Wei W, Cang J. 2024. Transformation of motion pattern selectivity from retina to superior colliculus. Journal of Neuroscience. DOI: 10.1523/JNEUROSCI.1704-23.2024, PMID: 38569924

Dhande OS, Huberman AD. 2014. Retinal ganglion cell maps in the brain: implications for visual processing. Current opinion in neurobiology 24:133–42. DOI: 10.1016/j.conb.2013.08.006, PMID: 24492089

Dipoppa M, Ranson A, Krumin M, Pachitariu M, Carandini M, Harris KD. 2018. Vision and Locomotion Shape the Interactions between Neuron Types in Mouse Visual Cortex. Neuron 98:602–615.e8. DOI: 10.1016/j.neuron.2018.03.037

Dräger UC, Hubel DH. 1975. Responses to visual stimulation and relationship between visual, auditory, and somatosensory inputs in mouse superior colliculus. Journal of neurophysiology 38:690–713. PMID: 1127462

Dreosti E, Odermatt B, Dorostkar MM, Lagnado L. 2009. A genetically encoded reporter of synaptic activity in vivo. Nature Methods 6:883–889. DOI: 10.1038/nmeth.1399

Edwards MA, Caviness Jr. VS, Schneider GE. 1986. Development of cell and fiber lamination in the mouse superior colliculus. Journal of Comparative Neurology 248:395–409. DOI: 10.1002/cne.902480308

Ellis EM, Gauvain G, Sivyer B, Murphy GJ. 2016. Shared and distinct retinal input to the mouse superior colliculus and dorsal lateral geniculate nucleus. Journal of Neurophysiology 116:602–610. DOI: 10.1152/jn.00227.2016, PMID: 27169509

Fabre JMJ, Van Beest EH, Peters AJ, Carandini M, Harris KD. 2023. Bombcell: automated curation and cell classification of spike-sorted electrophysiology data.

Feinberg EH, Meister M. 2014. Orientation columns in the mouse superior colliculus. Nature 519:229–232. DOI: 10.1038/nature14103

Franklin KBJ, Paxinos G. 2007. The Mouse Brain in Stereotaxic Coordinates, Third edition. Academic Press.

Hong YK, Kim I-J, Sanes JR. 2011. Stereotyped axonal arbors of retinal ganglion cell subsets in the mouse superior colliculus. The Journal of comparative neurology 519:1691–711. DOI: 10.1002/cne.22595, PMID: 21452242

Hoy JL, Farrow K. 2025. The superior colliculus. Current Biology 35:R164–R168. DOI: 10.1016/j.cub.2025.01.022, PMID: 40068606

Hubel DH, Wiesel TN. 1968. Receptive fields and functional architecture of monkey striate cortex. The Journal of physiology 195:215–43. PMID: 4966457

Hubel DH, Wiesel TN. 1962. Receptive fields, binocular interaction and functional architecture in the cat’s visual cortex. The Journal of physiology 160:106–54. PMID: 14449617

Huberman AD, Wei W, Elstrott J, Stafford BK, Feller MB, Barres BA. 2009. Genetic identification of an On-Off direction-selective retinal ganglion cell subtype reveals a layer-specific subcortical map of posterior motion. Neuron 62:327–34. DOI: 10.1016/j.neuron.2009.04.014, PMID: 19447089

Ibbotson M, Jung YJ. 2020. Origins of Functional Organization in the Visual Cortex. Frontiers in Systems Neuroscience 14. DOI: 10.3389/fnsys.2020.00010

Inayat S, Barchini J, Chen H, Feng L, Liu X, Cang J. 2015. Neurons in the Most Superficial Lamina of the Mouse Superior Colliculus Are Highly Selective for Stimulus Direction. Journal of Neuroscience 35:7992–8003. DOI: 10.1523/JNEUROSCI.0173-15.2015

International Brain Laboratory, Banga K, Boussard J, Chapuis GA, Faulkner M, Harris KD, Huntenburg JM, Hurwitz C, Lee HD, Paninski L, Rossant C, Roth N, Steinmetz NA, Windolf C, Winter O. 2022. Spike sorting pipeline for the International Brain Laboratory.

Ito S, Feldheim DA, Litke AM. 2017. Segregation of visual response properties in the mouse superior colliculus and their modulation during locomotion. The Journal of Neuroscience 37:3689–16. DOI: 10.1523/JNEUROSCI.3689-16.2017

Jang J, Song M, Paik SB. 2020. Retino-Cortical Mapping Ratio Predicts Columnar and Salt-and-Pepper Organization in Mammalian Visual Cortex. Cell Reports 30:3270–3279.e3. DOI: 10.1016/j.celrep.2020.02.038, PMID: 32160536

Jimenez LO, Tring E, Trachtenberg JT, Ringach DL. 2018. Local tuning biases in mouse primary visual cortex. Journal of Neurophysiology 120:274–280. DOI: 10.1152/JN.00150.2018/ASSET/IMAGES/LARGE/Z9K0061846420003.JPEG, PMID: 29668380

Jun JJ, Steinmetz NA, Siegle JH, Denman DJ, Bauza M, Barbarits B, Lee AK, Anastassiou CA, Andrei A, Aydın Ç, Barbic M, Blanche TJ, Bonin V, Couto J, Dutta B, Gratiy SL, Gutnisky DA, Häusser M, Karsh B, Ledochowitsch P, Lopez CM, Mitelut C, Musa S, Okun M, Pachitariu M, Putzeys J, Rich PD, Rossant C, Sun W, Svoboda K, Carandini M, Harris KD, Koch C, O’Keefe J, Harris TD. 2017. Fully integrated silicon probes for high-density recording of neural activity. Nature 551:232–236. DOI: 10.1038/nature24636

Kasai M, Isa T. 2021. Effects of light isoflurane anesthesia on organization of direction and orientation selectivity in the superficial layer of the mouse superior colliculus. Journal of Neuroscience JN-RM-1196–21. DOI: 10.1523/JNEUROSCI.1196-21.2021, PMID: 34872926

Kaschube M. 2014. Neural maps versus salt-and-pepper organization in visual cortex. Current Opinion in Neurobiology 24:95–102. DOI: 10.1016/j.conb.2013.08.017

Kay JN, Huerta ID la, Kim I-J, Zhang Y, Yamagata M, Chu MW, Meister M, Sanes JR. 2011. Retinal Ganglion Cells with Distinct Directional Preferences Differ in Molecular Identity, Structure, and Central Projections. Journal of Neuroscience 31:7753–7762. DOI: 10.1523/JNEUROSCI.0907-11.2011, PMID: 21613488

Kay RB, Triplett JW. 2017. Visual Neurons in the Superior Colliculus Innervated by Islet2+ or Islet2− Retinal Ganglion Cells Display Distinct Tuning Properties. Frontiers in Neural Circuits 11. DOI: 10.3389/fncir.2017.00073

Kerschensteiner D, Hardesty JF. 2022. Feature Detection by Retinal Ganglion Cells. Annual Review of Vision Science 8. DOI: 10.1146/ANNUREV-VISION-100419-112009

Kondo S, Yoshida T, Ohki K. 2016. Mixed functional microarchitectures for orientation selectivity in the mouse primary visual cortex. Nature Communications 7:13210. DOI: 10.1038/ncomms13210

Kovesi P. 2015. Good Colour Maps: How to Design Them. DOI: 10.48550/arXiv.1509.03700

Laniado DD, Maron Y, Gemmer JA, Sabbah S. 2025. A spherical code of retinal orientation selectivity enables decoding in ensembled and retinotopic operation. Cell Reports 44. DOI: 10.1016/j.celrep.2025.115373, PMID: 40023844

Lee KH, Tran A, Turan Z, Meister M. 2020. The sifting of visual information in the superior colliculus. eLife 9. DOI: 10.7554/eLife.50678

Li Y, Meister M. 2023. Functional cell types in the mouse superior colliculus. eLife 12:e82367. DOI: 10.7554/eLife.82367

Li Y, Turan Z, Meister M. 2020. Functional Architecture of Motion Direction in the Mouse Superior Colliculus. Current Biology 30:3304–3315.e4. DOI: 10.1016/j.cub.2020.06.023

Liang Y, Lu R, Borges K, Ji N. 2023. Stimulus edges induce orientation tuning in superior colliculus. Nature Communications 14:4756. DOI: 10.1038/s41467-023-40444-1

Liu X, Huang H, Snutch TP, Cao P, Wang L, Wang F. 2022. The Superior Colliculus: Cell Types, Connectivity, and Behavior. Neuroscience Bulletin 38:1519–1540. DOI: 10.1007/s12264-022-00858-1

Lopes G, Farrell K, Horrocks EA, Lee C-Y, Morimoto MM, Muzzu T, Papanikolaou A, Rodrigues FR, Wheatcroft T, Zucca S, Solomon SG, Saleem AB. 2021. Creating and controlling visual environments using BonVision. eLife 10:e65541. DOI: 10.7554/eLife.65541

Martin KAC, Schröder S. 2013. Functional heterogeneity in neighboring neurons of cat primary visual cortex in response to both artificial and natural stimuli. The Journal of Neuroscience 33:7325–44. DOI: 10.1523/JNEUROSCI.4071-12.2013, PMID: 23616540

Martin PR. 1986. The projection of different retinal ganglion cell classes to the dorsal lateral geniculate nucleus in the hooded rat. Experimental Brain Research 62:77–88. DOI: 10.1007/BF00237404

Mazade R, Alonso JM. 2017. Thalamocortical processing in vision. Visual Neuroscience 34:E007. DOI: 10.1017/S0952523817000049

Molotkov D, Ferrarese L, Boissonnet T, Asari H. 2023. Topographic axonal projection at single-cell precision supports local retinotopy in the mouse superior colliculus. Nature Communications 14:7418. DOI: 10.1038/s41467-023-43218-x

Najafian S, Koch E, Teh KL, Jin J, Rahimi-Nasrabadi H, Zaidi Q, Kremkow J, Alonso J-M. 2022. A theory of cortical map formation in the visual brain. Nature Communications 2022 13:1 **13**:1–20. DOI: 10.1038/s41467-022-29433-y

Nath A, Schwartz GW. 2016. Cardinal Orientation Selectivity Is Represented by Two Distinct Ganglion Cell Types in Mouse Retina. The Journal of Neuroscience 36:3208–21. DOI: 10.1523/JNEUROSCI.4554-15.2016, PMID: 26985031

Niell CM, Stryker MP. 2008. Highly selective receptive fields in mouse visual cortex. The Journal of Neuroscience 28:7520–36. DOI: 10.1523/JNEUROSCI.0623-08.2008, PMID: 18650330

Ohki K, Chung S, Ch’ng YH, Kara P, Reid RC. 2005. Functional imaging with cellular resolution reveals precise micro-architecture in visual cortex. Nature 433:597–603. DOI: 10.1038/nature03274, PMID: 15660108

Oommen BS, Stahl JS. 2008. Eye orientation during static tilts and its relationship to spontaneous head pitch in the laboratory mouse. Brain Research 1193:57–66. DOI: 10.1016/j.brainres.2007.11.053

Pachitariu M, Sridhar S, Pennington J, Stringer C. 2024. Spike sorting with Kilosort4. Nature Methods 21:914–921. DOI: 10.1038/s41592-024-02232-7

Pachitariu M, Stringer C, Schröder S, Dipoppa M, Rossi LF, Carandini M, Harris KD. 2016. Suite2p: beyond 10,000 neurons with standard two-photon microscopy. bioRxiv 061507. DOI: 10.1101/061507

Pologruto TA, Sabatini BL, Svoboda K. 2003. ScanImage: flexible software for operating laser scanning microscopes. Biomedical Engineering Online 2:13. DOI: 10.1186/1475-925X-2-13, PMID: 12801419

Ringach DL, Mineault PJ, Tring E, Olivas ND, Garcia-Junco-Clemente P, Trachtenberg JT. 2016. Spatial clustering of tuning in mouse primary visual cortex. Nature Communications 7:12270. DOI: 10.1038/ncomms12270

Roth ZN, Heeger DJ, Merriam EP. 2018. Stimulus vignetting and orientation selectivity in human visual cortex. eLife 7. DOI: 10.7554/eLife.37241

Sabbah S, Gemmer JA, Bhatia-Lin A, Manoff G, Castro G, Siegel JK, Jeffery N, Berson DM. 2017. A retinal code for motion along the gravitational and body axes. Nature 546:492–497. DOI: 10.1038/nature22818

Savier EL, Chen H, Cang J. 2019. Effects of Locomotion on Visual Responses in the Mouse Superior Colliculus. The Journal of Neuroscience 1854–19. DOI: 10.1523/JNEUROSCI.1854-19.2019, PMID: 31570535

Schmidt KE, Wolf F. 2021. Punctuated evolution of visual cortical circuits? Evidence from the large rodent Dasyprocta leporina, and the tiny primate Microcebus murinus. Current Opinion in Neurobiology 71:110–118. DOI: 10.1016/J.CONB.2021.10.007, PMID: 34823047

Schröder S, Steinmetz NA, Krumin M, Pachitariu M, Rizzi M, Lagnado L, Harris KD, Carandini M. 2020. Arousal Modulates Retinal Output. Neuron 107:487–495.e9. DOI: 10.1016/j.neuron.2020.04.026

Shi X, Barchini J, Ledesma HA, Koren D, Jin Y, Liu X, Wei W, Cang J. 2017. Retinal origin of direction selectivity in the superior colliculus. Nature Neuroscience 20:550–558. DOI: 10.1038/nn.4498

Sibille J, Gehr C, Benichov JI, Balasubramanian H, Teh KL, Lupashina T, Vallentin D, Kremkow J. 2022. High-density electrode recordings reveal strong and specific connections between retinal ganglion cells and midbrain neurons. Nature Communications 13:5218. DOI: 10.1038/s41467-022-32775-2

Stitt I, Galindo-Leon E, Pieper F, Engler G, Engel AK. 2013. Laminar profile of visual response properties in ferret superior colliculus. Journal of neurophysiology 110:1333–45. DOI: 10.1152/jn.00957.2012, PMID: 23803328

Tiriac A, Bistrong K, Pitcher MN, Tworig JM, Feller MB. 2022. The influence of spontaneous and visual activity on the development of direction selectivity maps in mouse retina. Cell Reports 38. DOI: 10.1016/j.celrep.2021.110225, PMID: 35021080

Vita DJ, Orsi FS, Stanko NG, Clark NA, Tiriac A. 2024. Development and organization of the retinal orientation selectivity map. Nature Communications 15:4829. DOI: 10.1038/s41467-024-49206-z

Wang L, Sarnaik R, Rangarajan K, Liu X, Cang J. 2010. Visual receptive field properties of neurons in the superficial superior colliculus of the mouse. The Journal of Neuroscience 30:16573–84. DOI: 10.1523/JNEUROSCI.3305-10.2010, PMID: 21147997

Warren RA, Zhang Q, Hoffman JR, Li EY, Hong YK, Bruno RM, Sawtell NB. 2021. A rapid whisker-based decision underlying skilled locomotion in mice. eLife 10:e63596. DOI: 10.7554/eLife.63596

Wheatcroft T, Saleem AB, Solomon SG. 2022. Functional Organisation of the Mouse Superior Colliculus. Frontiers in Neural Circuits 16:792959. DOI: 10.3389/fncir.2022.792959, PMID: 35601532

Yaakov H, Heukamp AS, Riccitelli S, Rivlin-Etzion M. 2025. Differential topographic organization and retinal inheritance of direction and orientation selectivity in the visual thalamus. Nature Communications 16:9303. DOI: 10.1038/s41467-025-64321-1

Yen S-C, Baker J, Gray CM. 2007. Heterogeneity in the responses of adjacent neurons to natural stimuli in cat striate cortex. Journal of neurophysiology 97:1326–41. DOI: 10.1152/jn.00747.2006, PMID: 17079343

Yu P, Yang Y, Gozel O, Oldenburg I, Dipoppa M, Rossi LF, Miller K, Adesnik H, Ji N, Doiron B. 2025. Circuit-Based Understanding of Fine Spatial Scale Clustering of Orientation Tuning in Mouse Visual Cortex. bioRxiv 2025.02.11.637768. DOI: 10.1101/2025.02.11.637768, PMID: 39990306

Zhao X, Liu M, Cang J. 2014. Visual Cortex Modulates the Magnitude but Not the Selectivity of Looming-Evoked Responses in the Superior Colliculus of Awake Mice. Neuron 84:202–213. DOI: 10.1016/j.neuron.2014.08.037

